# Distinct genome architecture underlies fine-scale population differentiation in two common European bumblebees (*Bombus pascuorum* and *Bombus lapidarius*)

**DOI:** 10.1101/2024.05.09.593344

**Authors:** Lauren Cobb, Markus A. K. Sydenham, Anders Nielsen, Bastiaan Star

## Abstract

Bumblebees are keystone pollinators which facilitate the reproduction of a wide range of wild and agricultural plants. Their abundance and diversity have been severely reduced by anthropogenic stressors such as land-use change and widespread habitat fragmentation. However, we lack a comprehensive understanding of bumblebee population structure and local adaptation in response to human-altered landscapes. We here discover surprisingly fine-scaled population structure (e.g. ∼300km) within two widely occurring bumblebee species, *Bombus lapidarius* and *Bombus pascuorum*, by analysing whole genome data of 106 specimens from 7 sites in Northern Europe. Our sample range encompasses a mosaic of land-use types with varying levels of habitat fragmentation and natural oceanic barriers. While the observed population structure is largely associated with reduced gene flow across natural barriers, we also detect significant divergence between populations sampled from more fragmented, agricultural landscapes. Furthermore, we identify species-specific patterns of population structure which are underpinned by distinct genomic architecture. Whereas genetic divergence in *B. lapidarius* is spread relatively evenly across the genome, divergence in *B. pascuorum* is concentrated within several megabase-sized genomic regions with significantly elevated differentiation – including a putative chromosomal inversion – which may underlie well-known colour polymorphisms across its range. Our observations reveal unexpectedly high levels of inter- and intraspecific genomic diversity within the bumblebee genus, and highlight the necessity of increasing our understanding of bumblebee population structure and connectivity to design optimal bumblebee conservation strategies.

**Significance statement:** Anthropogenic stressors such as habitat fragmentation have severe impacts on bumblebee abundance and diversity, yet little is known about how bumblebee populations are structured in human-altered landscapes. We analyse whole-genome data from two common bumblebee species (*Bombus lapidarius* and *Bombus pascuorum*) across Northern Europe to uncover species-specific patterns of spatial population differentiation and local adaptation, including a chromosomal rearrangement in *B. pascuorum*. Importantly, our results imply that many of the fragmented bumblebee habitats in Europe comprise locally distinct populations with limited gene flow in between. These findings are therefore of major importance for our overall understanding of bumblebee genomic variation, connectivity and adaptation, offering fundamental insights that are required to effectively mitigate the effects of human activities on wild bee biodiversity.

## Introduction

Bumblebees are keystone temperate pollinators (1): they are abundant, primarily generalist foragers, with wide-ranging distributions encompassing various different temperate ecosystems. These ecological features allow them to facilitate the reproduction of a diverse array of wild and agricultural plant species, many of which are pollinated predominantly by bumblebees (2, 3). Over recent decades, however, evidence for an alarming global trend of insect pollinator loss has accumulated (4–6). Bumblebees are no exception, with population declines and local extinctions reported throughout Europe, North America and Asia (2, 7–10). If such trends persist, these widespread losses in bumblebee abundance and diversity have potentially catastrophic implications, for both the biodiversity of wild temperate plants which underpin a wide range of ecological functions, and for the preservation of agricultural food security (4, 11).

The root causes of bumblebee declines are complex, and are thought to be the result of a combination of anthropogenic stressors (8, 12, 13). Alongside human-introduced parasites and pathogens, climate change and agrochemical exposure, human-driven land-use change is widely considered a major driver (8, 14–16). Alteration of landscapes through anthropogenic activities, such as urbanisation and agricultural intensification, severely depletes bumblebee foraging and nesting resources (10, 14). Bumblebee habitats therefore become heavily fragmented, reducing habitat connectivity throughout their range, and, as a result, bumblebee dispersal and gene flow is obstructed (17, 18). We are yet to reach a comprehensive understanding of the mechanisms linking these environmental stressors to bumblebee declines, particularly given that such trends are not universal across bumblebee species – while some are profoundly affected by these threats, others appear much more resilient (19). In order to deliver effective conservation strategies to mitigate bumblebee declines and their associated environmental consequences, it is essential that we increase our knowledge of the impacts of anthropogenic land-use on the various interacting facets of bumblebee biology, from the ecological level to the genomic level.

Habitat fragmentation and reduced population connectivity can have devastating biological consequences: small, isolated populations with limited gene flow suffer from loss of genetic diversity due to genetic drift and increased inbreeding, resulting in reduced individual fitness and limited ability to adapt to environmental change (20, 21). These genetic consequences may be particularly impactful for bumblebees due to several biological traits. Bumblebees are haplodiploid, meaning fewer gene copies are present in the population in comparison to diploid species, and have a eusocial breeding structure, therefore each nest can represent few breeding pairs despite consisting of many individuals. Both characteristics reduce their effective population size (*Ne*) compared to diploid, non-eusocial species with comparable abundance (22, 23). In addition, bumblebees have single-locus complementary sex determination: females develop from fertilised eggs which are heterozygous at the sex-determining locus, and males develop from unfertilised eggs (24, 25). However, homozygosity at the sex-determining locus, made more likely by small populations and inbreeding, results in the development of the genetic ‘dead end’ of a sterile diploid male (25–27). Combined, these biological characteristics compound the risk of inbreeding depression and local extinction in bumblebees.

Despite these increased genetic vulnerabilities, studies of bumblebee population structure and genetic diversity have largely been performed using limited numbers of molecular markers (24, 25, 27–35). The results of these studies vary between species, with both declining and common species exhibiting either significant (25, 27, 29, 35) or minimal (29, 33) genetic structure across a range of spatial scales. While the use of few molecular markers is relatively robust for the detection of strong population structure, such approaches lack the sensitivity, accuracy and resolution of whole genome analyses (36–38). For instance, reductions in effective population size and signs of inbreeding in the declining North American bumblebee *Bombus terricola* were identified by partial genome sequencing (39), while significant population structure and signs of recent bumblebee adaptation to anthropogenic threats have been detected through RADSeq (40). While no evidence of genomic barriers was detected across the largely undisturbed northern Scandinavian range of several montane bumblebee species (e.g. *Bombus lapponicus*, *Bombus monticola*), significant structure was found in *Bombus pascuorum* using whole genome sequencing (41). Nonetheless, given the scarcity of literature surrounding this subject, the study of bumblebee genomic structure and adaptation in response to altered and fragmented landscapes is in urgent need of increased scientific attention.

Here, we conduct a comparative population genomics study of two common bumblebee species, *Bombus pascuorum* and *Bombus lapidarius.* Both species are among the most widespread and abundant bumblebees in temperate Europe (3), and are important pollinators of a range of wild and agricultural plants (2). *Bombus pascuorum* (the common carder bee), a medium-tongue-length forager with a preference for agricultural landscapes (2, 42), is the most polytypic of all bumblebees, exhibiting over 20 colour morphs across its expansive European distribution (43), which indicates a significant degree of geographic divergence (44). *Bombus lapidarius* (the red-tailed bumblebee) is shorter-tongued and commonly inhabits semi-natural, managed and urban habitats, suggesting a relative resilience to anthropogenic land*-*use change in comparison to other, declining species (45). Indeed, the past century has seen *B. lapidarius* become increasingly dominant in northern European agricultural landscapes (46). While neither species is of immediate conservation concern, as pollinators of a wide range of plants it essential that they are nonetheless included in conservation research. Moreover, it has been predicted that over one third of bumblebee species currently classified as ‘Least Concern’ will lose at least 30% of their habitat to anthropogenic land-use change in the coming decades (47). Significant genetic structure has previously been identified in both species (29), but with the use of a limited number of genetic markers, and solely at a continental scale: the sample sites between which structure was found were 1650 km apart, with no evidence of structure at shorter distances. Whole genome analyses of *Bombus pascuorum* throughout Sweden detected substructure at considerably finer spatial scales, associated with island populations and differences in colour morphology, although the genetic architecture underlying this differentiation was not further explored (41). We therefore still have a limited understanding of the drivers of bumblebee population structure and adaptation at finer spatial resolutions.

We address this knowledge gap by using whole-genome sequencing to investigate the population structure of *Bombus pascuorum* and *Bombus lapidarius* throughout southern Norway, southern Sweden, Denmark and North Germany. Our sample range is home to four *B. pascuorum* morphs: *B. p. sparreanus* in western Norway, *B. p. pallidofacies* in south-eastern Norway and southern Sweden, *B. p. floralis* in Germany and most of Denmark, and *B. p. mniorum* in Zealand, Denmark (43, 48). This range also incorporates a gradient of habitat fragmentation, with higher rates of human land-use present in Germany and Denmark than in Norway and Sweden, where landscapes are typically more heterogenous (49). Indeed, parts of Norway and Sweden have been identified by models as potential refugia for European bumblebees in the face of ongoing human-driven habitat loss (47). In addition, our sample range is currently unstudied, and therefore fills a crucial geographical blind spot in the field of bumblebee population genomics.

## Results

### Whole Genome Sequencing

We analysed 5,603,463,226 reads, ranging from 261,576 to 96,527,020 per specimen, with a mean coverage of 16X (**Supplementary Table 1**, **2**). Following individual and quality filtering, we analyse ∼1.5 million SNPs in 45 *B. lapidarius* specimens and ∼2.8 million SNPs in 61 *B. pascuorum* specimens with a mean coverage of 21X.

### Genomic structure and differentiation

Principal Component Analysis (PCA) of both *B. lapidarius* and *B. pascuorum* whole-genome datasets indicate several distinct clusters across each sample range (**Figure 1**). For *B. lapidarius*, the first two PCs exhibit significant substructure (Tracy-Widom statistics - PC1: twstat = 40.31, p < 0.01, eigenvalue = 2.16; PC2: twstat = 13.04, p < 0.01, eigenvalue = 1.12) and samples fall into 3 main clusters: Norwegian/Swedish samples, North Jutland/Funen samples, and German samples. The two Zealand samples fall close to the Norway/Sweden cluster on PC1, but segregate in the direction of the remaining Danish samples along PC2 (**Figure 1b**). For *B. pascuorum,* the first three PCs exhibit significant substructure (Tracy-Widom statistics - PC1: twstat = 58.91, p < 0.01, eigenvalue = 2.06; PC2: twstat = 7.03, p < 0.01, eigenvalue = 1.18; PC3: twstat = 3.002, p < 0.01, eigenvalue = 1.15), and samples fall into several clusters. Norwegian and Swedish samples form one cluster separated from Danish and German samples, which are each distinctly grouped along PC2 (**Figure 1d**). Furthermore, the North Jutland and North Germany clusters are strongly differentiated along PC3 **(Supplementary Figure 1)**.

**Figure 1.**
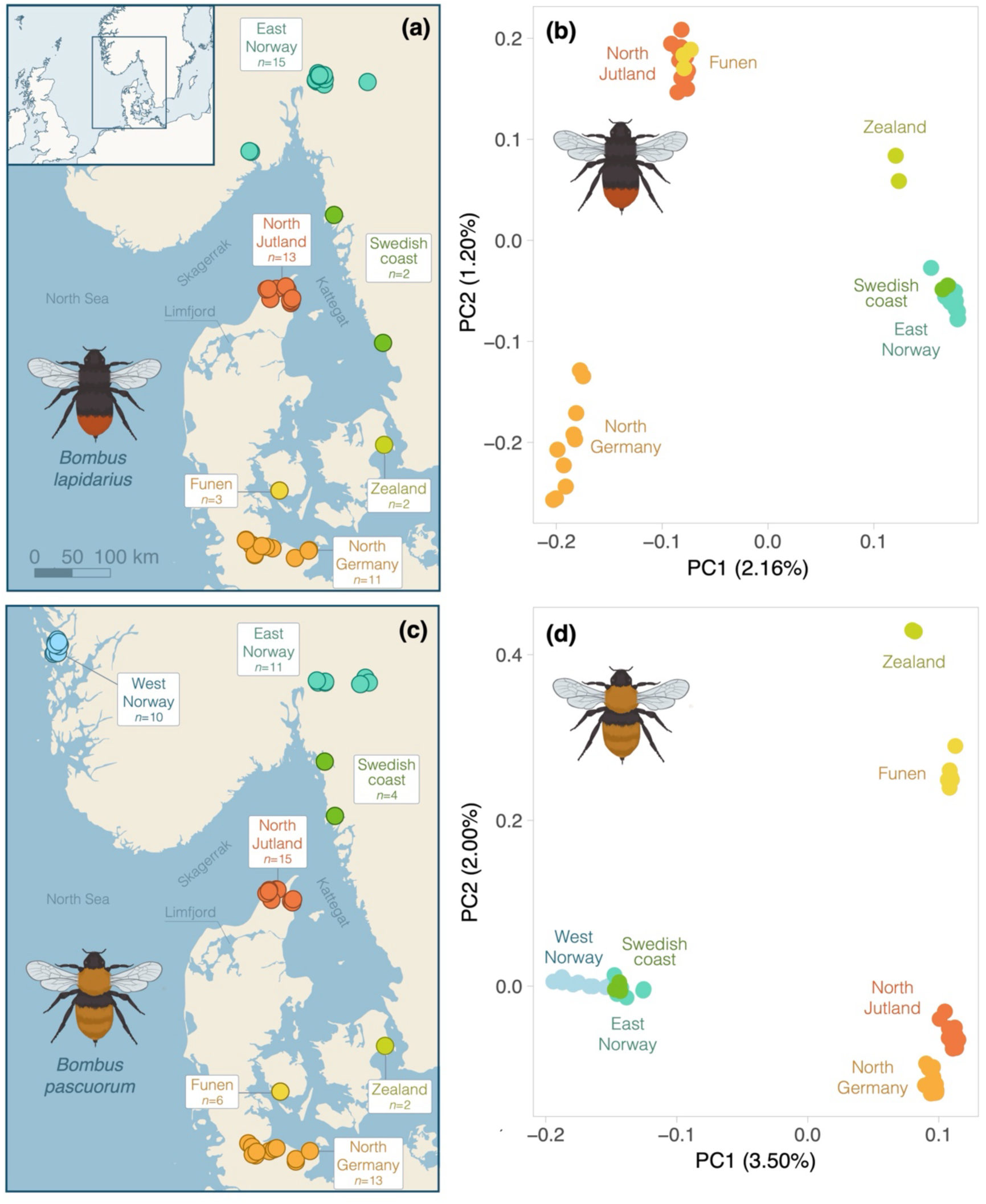
Distinct genome-wide population structure in *Bombus lapidarius* (*n=*45) and *Bombus pascuorum* (*n*=61) across southern Norway, southern Sweden, Denmark and north Germany. **(a & c)** Maps representing the sampling locations (coloured dots) of all *B. lapidarius* and *B. pascuorum* samples, respectively. Points represent individual sampling locations, and colours indicate sample site assignments. Labels indicate the number of individuals sampled within each site. **(b & d)** Principal component analyses (PCA) of **(b)** 698,786 SNPs from 45 *B. lapidarius* individuals and **(d)** 1,863,457 SNPs from 61 *B. pascuorum* individuals. Points represent individual samples and colours indicate sample site assignments. Percentages indicate the proportion of genomic variation explained by each PC axis.

These results are concurrent with those of the model-based clustering analysis (ADMIXTURE) (**Figure 2**). For *B. lapidarius*, German and Norwegian/Swedish samples are strongly distinct from each other for K=2, K=3 and K=4. For K=2, the North Jutland and Funen samples cluster predominantly with the German samples, and the Zealand samples cluster with the Norwegian/Swedish samples. For K=3 and K=4, North Jutland and Funen samples form one cluster, with some admixture from both the German and Norwegian/Swedish ancestral populations, while the Zealand samples cluster with the Norwegian/Swedish samples, with slight admixture from the Danish ancestral population. Finer substructure within the Danish samples is revealed by K=4 (**Figure 2a**). Cross-validation error is lowest for K=1, increases slightly for K=2, followed by steeper increases for each greater value of K (**Supplementary Figure 2a**). Cluster-based model validation using evalAdmix however indicates a good model fit for each K value, increasing slightly with increased K (**Supplementary Figure 3a**).

**Figure 2.**
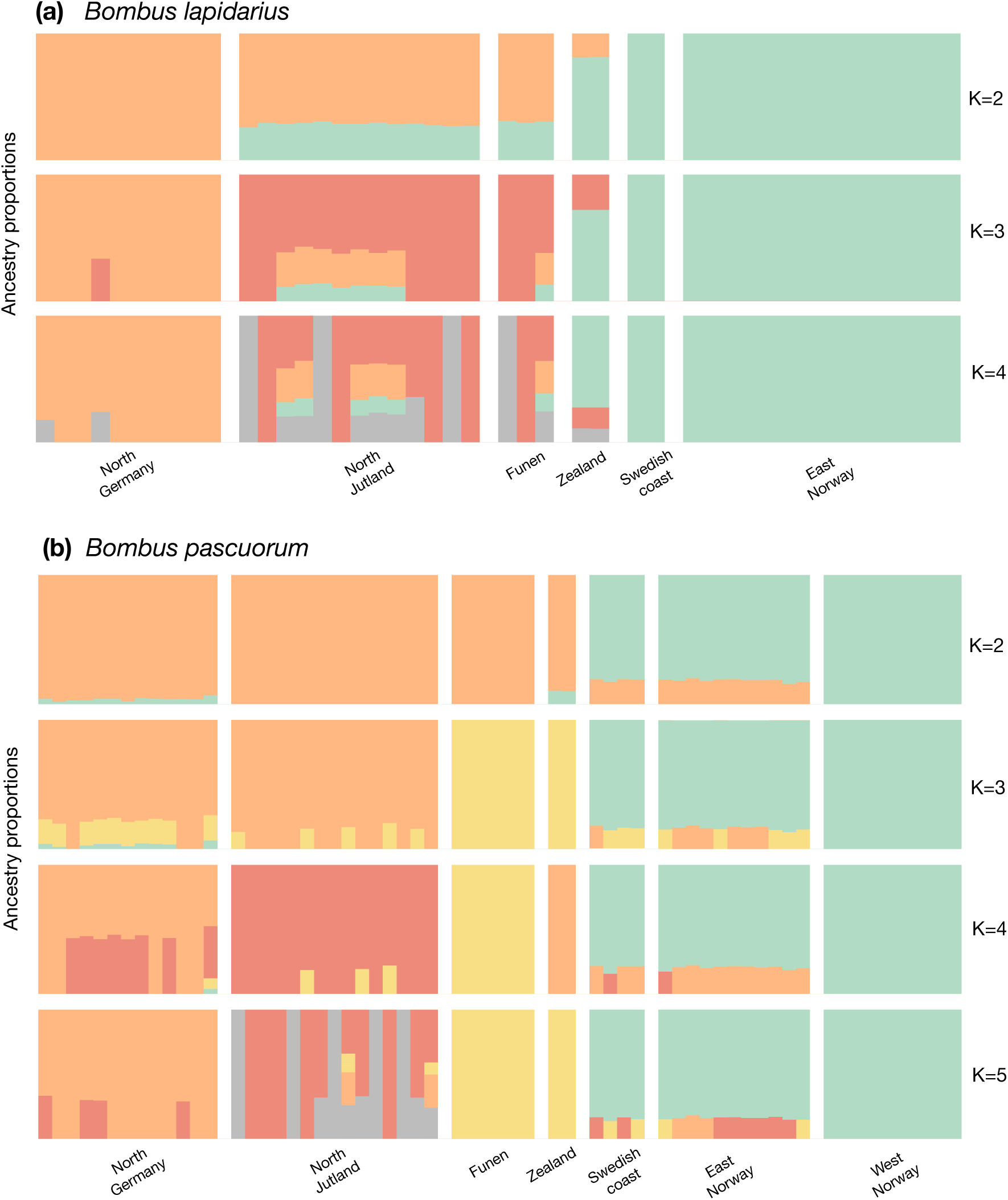
Population structure and admixture of *B lapidarius* (*n=*45) and *B. pascuorum* (*n*=61). Admixture plots showing the ancestry proportions of the best values of K for (a) *Bombus lapidarius* and (b) *Bombus pascuorum* sampled across southern Norway, southern Sweden, Denmark and north Germany. Each column represents an individual, and each colour represents a different hypothetical ancestral (K) population to which a proportion of ancestry is assigned per individual. Individuals are grouped by sampling site, which are separated by vertical white lines, and different values of K are separated by horizontal white lines. **(a)** Ancestry proportions were estimated from ADMIXTURE analysis of 698,786 SNPs for K=2 - K=4. **(b)** Ancestry proportions were estimated from ADMIXTURE analysis of 1,863,457 SNPs for K=2 - K=5.

For *B. pascuorum*, the Norwegian/Swedish samples and the German/Danish samples form two distinct clusters for K=2, with a small amount of admixture exhibited by the German, Zealand, Swedish, and Norwegian samples. For K=3, Funen and Zealand samples form one cluster. For K=4, the German and North Jutland populations segregate, with most of the German samples retaining considerable admixture from the North Jutland ancestral population. In addition, the two Zealand samples cluster with the North German samples. For K=5, Zealand samples cluster with Funen samples, and North Jutland samples exhibit some finer substructure (**Figure 2b**). Cross-validation error is lowest for K=1, increases slightly for K=2, followed by steeper increases for each greater value of K (**Supplementary Figure 2b**). Cluster-based model validation using evalAdmix however indicates a good model fit for each K value, increasing slightly with increased K (**Supplementary Figure 3b**).

Relative population differentiation for *B. lapidarius*, as measured by pairwise Hudson’s Fst values between the clusters inferred by the PCA and ADMIXTURE analyses, is greatest between the Norway/Sweden cluster and the two remaining clusters. The highest differentiation is that of the Norway/Sweden cluster from the German cluster, which is almost 2X greater than its differentiation from the Jutland/Funen cluster, and approximately 4X greater than the differentiation between the Germany and Jutland/Funen clusters (**Table 1**). Relative population differentiation for *B. pascuorum* is also greatest between the Norway/Sweden cluster and all remaining clusters. The highest differentiation is that of the Norway/Sweden cluster from the Funen cluster, closely followed by its differentiation from the German cluster, both of which are between 3X and 4X greater than the differentiation found between all remaining clusters (Table 2). For both species, absolute population differentiation, as measured by Dxy, did not differ significantly between the sites (**Table 1**; **Table 2**).

**Table 1.** Between-population pairwise Hudson’s Fst values (lower triangle), pairwise dxy values (upper triangle), and Tajima’s D (center diagonal) for 43 *Bombus lapidarius* individuals.

|  | Germany | Jutland/<br>Funen | Norway/<br>Sweden |
| --- | --- | --- | --- |
| Germany<br>(n=10) | -0.37 | 0.004 | 0.004 |
| Jutland/<br>Funen<br>(n=16) | 0.010 | -0.21 | 0.004 |
| Norway/<br>Sweden<br>(n=17) | 0.045 | 0.026 | 0.22 |

**Table 2.** Between-population pairwise Hudson’s Fst values (lower triangle), pairwise dxy values (upper triangle), and Tajima’s D (center diagonal) for 59 *Bombus pascuorum* individuals.

|  | Germany | Funen | Jutland | Norway/<br>Sweden |
| --- | --- | --- | --- | --- |
| Germany<br>(n=13) | -0.61 | 0.009 | 0.009 | 0.009 |
| Funen<br>(n=6) | 0.007 | -0.31 | 0.009 | 0.009 |
| Jutland<br>(n=15) | 0.008 | 0.010 | -0.53 | 0.009 |
| Norway/<br>Sweden<br>(n=25) | 0.036 | 0.037 | 0.029 | -0.51 |

### Genomic diversity and inbreeding

For *B. lapidarius,* Tajima’s D is between 1 and -1 for all clusters, with Germany and Jutland/Funen clusters exhibiting negative values and the Norway/Sweden cluster exhibiting a positive value (**Table 1**). Inbreeding values as measured by F statistics are low, ranging between -0.03 and 0.04, with no statistically significant difference between populations (**Supplementary Table 3**). Levels of heterozygosity differed significantly between the sites (F = 20.2, df = 4, p < 0.001; mean: 30.2±0.4). The East Norway/Sweden cluster exhibits significantly lower heterozygosity than the Funen (adjusted p < 0.001), North Germany (adjusted p < 0.001) and North Jutland populations (adjusted p < 0.001). The Zealand individuals also exhibit significantly lower heterozygosity than the Funen (adjusted p < 0.05) and North Jutland (adjusted p < 0.05) populations (**Figure 3a**). 10372 RoHs were detected from *B. lapidarius* genomes, ranging up to 382kb in length and including 255 RoHs longer than 100kb (mean length: 26.0kb±24.5). RoH length was initially found to be statistically different between populations (Kruskal-Wallis chi-squared = 11.40, df = 4, p < 0.05), However, a post-hoc Dunn test produced no significant adjusted p values (**Figure 3b)**. Percentage of the genome occupied by RoHs mostly ranges between <0.1% and 3%, with a mean of 1.69±0.77, and was found to be statistically different between populations (F = 3.94, df = 4, p < 0.01). A post-hoc Tukey Honest Significant Difference test showed that Germany and Norway/Sweden were the only significantly different pair of populations (adjusted p < 0.01) (**Figure 3c**).

**Figure 3.**
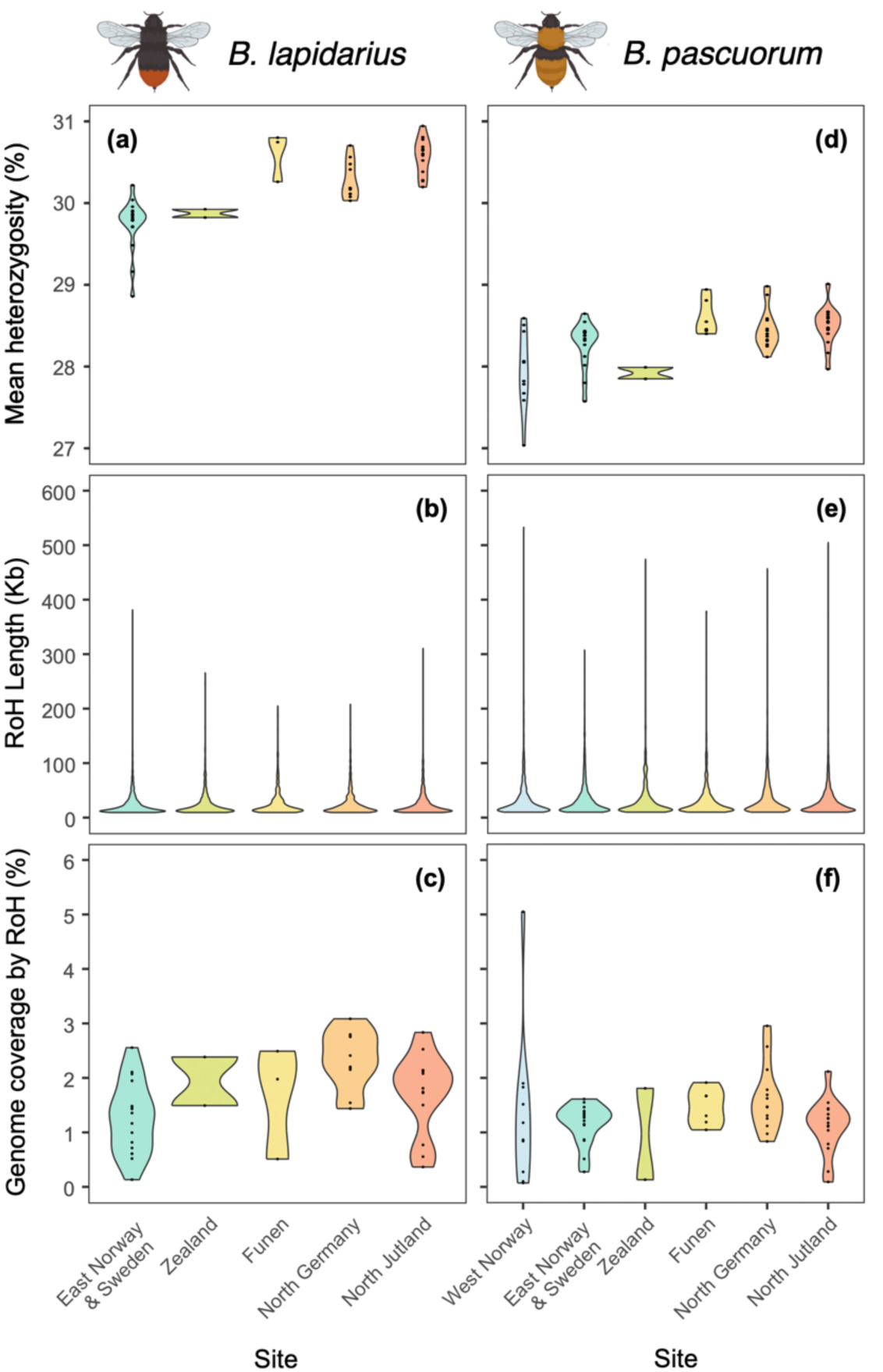
Distributions of mean percentage heterozygosity (a & d), RoH length (b & e; minimum length: 10kb), and percentage of the genome covered by RoHs (c & f) of 61 *Bombus pascuorum* individuals and 45 *Bombus lapidarius* individuals. *B. pascuorum* individuals were sampled across West Norway (*n*=10), East Norway & Sweden (*n*=15), Zealand (*n*=2), Funen (*n*=6), North Germany (*n*=13) and North Jutland (*n*=15), and analyses were generated from 2,839,545 SNPs. *B. lapidarius* individuals were sampled across North Germany (*n*=10), North Jutland (*n*=13), Funen (*n*=3), Zealand (*n*=2), the Swedish coast (*n*=2) and Eastern Norway (*n*=15), and analyses were generated from 1,464,934 SNPs.

For *B. pascuorum*, Tajima’s D is between 0 and -1 for all clusters (**Table 2**). Inbreeding values as measured by F statistics are low, ranging between -0.03 and 0.04, with no statistically significant difference between populations (**Supplementary Table 4).** The West Norway population exhibits significantly lower levels of heterozygosity than the Funen (adjusted p < 0.01), North German (adjusted p < 0.01) and North Jutland (adjusted p < 0.001) populations (F = 6.33, df =5, p < 0.001). No other significant differences in heterozygosity between populations were found (mean: 28.3±0.4; **Figure 3d**). 7650 RoHs were detected from *B. pascuorum* genomes, ranging up to 533kb in length and including 321 RoHs longer than 100kb (mean length: 32.0kb±33.0). RoH length was initially found to be statistically different between populations (*B. pascuorum*: Kruskal-Wallis chi-squared = 14.12, df = 5, p-value < 0.05; *B. lapidarius:* Kruskal-Wallis chi-squared = 11.40, df = 4, p < 0.05). However, a post-hoc Dunn test produced no significant adjusted p values (**Figure 3e)**. Percentage of the genome occupied by RoHs mostly ranges between <0.1% and 3%, except one sample from West Norway with a value of 5%, with a mean of 1.30±0.75 and no significant differences between populations (**Figure 3f**). *B. lapidarius* samples exhibit significantly greater mean percentage heterozygosity than *B. pascuorum* (t = 22.44, df = 82.37, p < 0.001). *B. lapidarius* RoHs are significantly shorter than those of *B. pascuorum* (W = 34571598, p < 0.01), and RoHs occupy a significantly greater mean percentage of the *B. lapidarius* genome than that of *B. pascuorum* (t = 2.60, df = 93.521, p < 0.05).

### Genome-wide divergence and genomic architecture

For *B. lapidarius*, pairwise genome-wide divergence between the clusters is low and distributed fairly uniformly among the first 95 scaffolds of the contig-level genome assembly (**Figure 4a**). Fst rarely exceeds background levels of divergence of approximately 0.01 for the Germany and Jutland/Funen pair, 0.04 for the Germany and Norway/Sweden pair, and 0.02 for the Jutland/Funen and Norway/Sweden pair. Several narrow outlying regions occur in all pairwise comparisons with Fst values that fall above the 99^th^ percentile distribution. The strongest and most frequent of these occur between the German and Norway/Sweden pair, and the fewest occur between the Germany and North Jutland/Funen pair. The Germany and Norway/Sweden pairwise comparison shares regions of elevated divergence on scaffold 1 and scaffold 20 with the Germany and Jutland/Funen pair, and regions of elevated divergence on scaffolds 8, 25, 46, 61 and 68 with the Jutland/Funen and Norway/Sweden pair. None of these regions showed distinct bi-allelic inheritance patterns (**Supplementary Figures 4-11**). The Germany and Norway/Sweden pairwise comparison also exhibits a unique region of elevated divergence on scaffold 33, comprising 0.1 Mbp and 134 SNPs. PCA of this region separates samples into three clusters along PC1, which explains 20% of overall variation. Percentage heterozygosity of these three clusters suggest a biallelic genotype pattern, with Cluster 1 and Cluster 3 exhibiting lower percentage heterozygosity than Cluster 2 (Cluster 1 mean: 10.68±5.25; Cluster 2 mean: 43.64±6.83; Cluster 3 mean: 25.55±6.28; **Supplementary Figure 12**).

**Figure 4.**
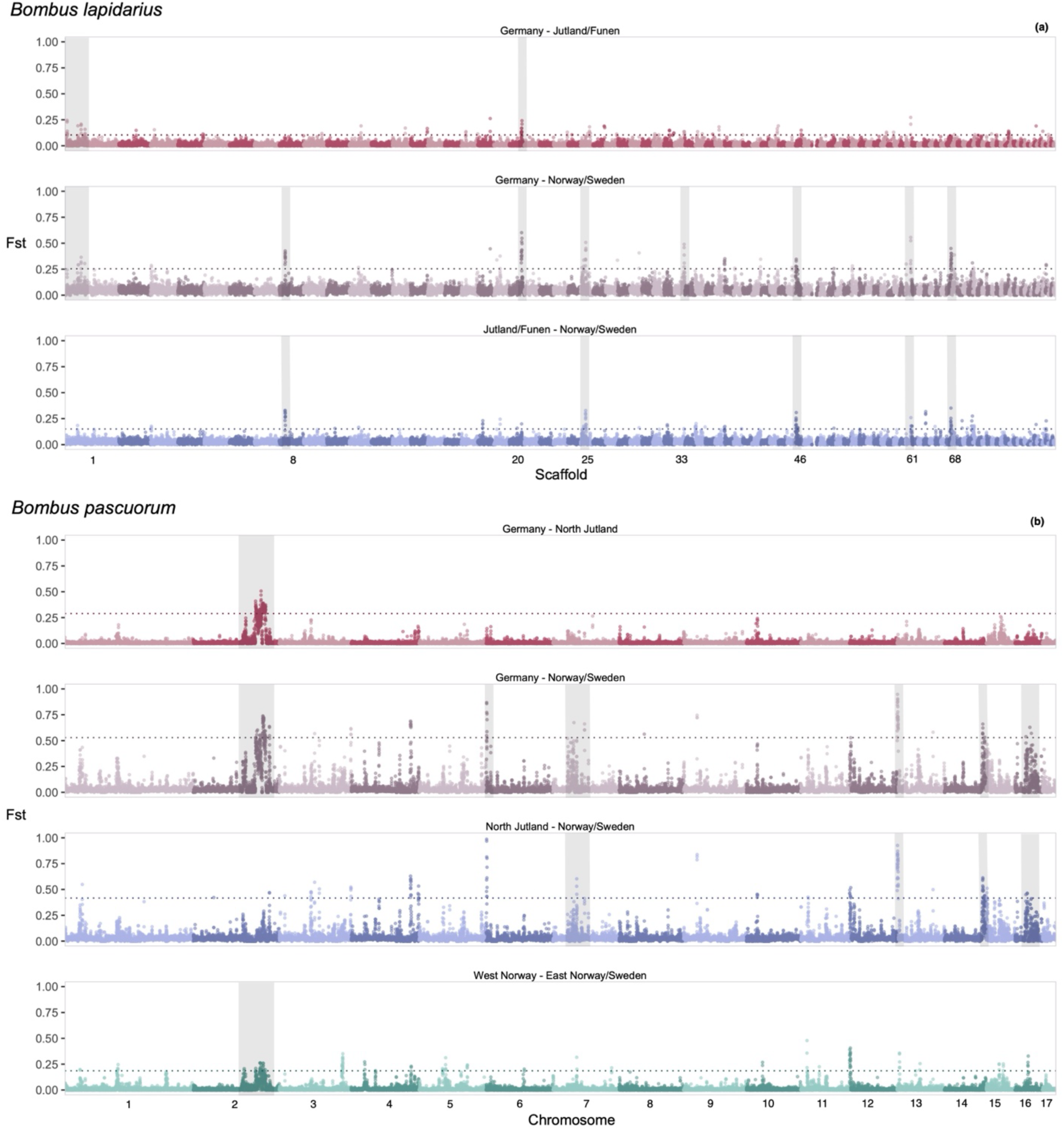
Distinct pairwise genomic differentiation (Fst) landscapes between (a) three *Bombus lapidarius* populations and (b) four *Bombus pascuorum* populations. Fst was calculated in sliding windows across the genome with a window size 25kb and a step size of 12.5kb. Red, grey and blue graphs indicate different population pairs. Light and dark versions of each colour signify alternating consecutive scaffolds (*B. lapidarius*) or chromosomes (*B. pascuorum*). Dotted lines represent the 99^th^ percentile of each Fst frequency distribution. Shaded boxes indicate peaks of elevated divergence which are subject to further local analyses. **(a)** The three *B. lapidarius* populations are North Jutland/Funen (*n*=16), Germany (*n*=10) and Norway/Sweden (*n*=17). Fst was calculated across the first 95 scaffolds of the contig-level genome, comprising 1,390,497 SNPs. **(b)** The four *B. pascuorum* populations are North Jutland (*n*=15), Germany (*n*=13), Norway/Sweden (*n*=25) and West Norway (*n*=10). Fst was calculated across the 17 assembled chromosomes of the genome, comprising 2,784,282 SNPs.

For *B. pascuorum*, we identify considerably broader genomic regions with elevated Fst divergence above the 99^th^ percentile distribution for all four pairwise comparisons between the clusters (**Figure 4b**). The Germany and Norway/Sweden pair exhibit the strongest and most frequent elevated peaks throughout the genome. The Germany and North Jutland pair has fewer elevated peaks, with one region of very strong divergence across several Mbps (million base pairs) on the second chromosome peaking at approximately 0.4 Fst. This region is also observed between the Germany and Norway/Sweden clusters, peaking at approximately 0.7 Fst, and between the West Norway and East Norway/Sweden clusters, peaking at approximately 0.25 Fst.

This highly divergent region occurs between 16.9-18.6 Mbp and 18.85-19.74 Mbp on chromosome 2, and combined contains 23,748 SNPs (**Figure 5a**). PCA of these regions separates all samples into three distinct clusters along PC1, which explains 44% of overall variation (**Figure 5b**). Percentage heterozygosity of these three clusters indicate a clear biallelic genotype pattern, with Cluster 1 and Cluster 3 exhibiting very low percentage heterozygosity (Cluster 1 mean: 10.93±1.46; Cluster 3 mean: 18.32±4.89) and Cluster 2 exhibiting much greater percentage heterozygosity (mean: 55.37±6.92; **Figure 5c**). Moreover, a clear geographical trend is present within this genotype pattern. Norwegian and Swedish samples occur in either Cluster 1 or Cluster 2. Most North Jutland samples occur in either Cluster 1 or 2, with one sample in Cluster 3. Funen samples occur in either Cluster 2 or Cluster 3. Most North German samples occur in Cluster 3, with two samples in Cluster 1 and two in Cluster 2.

**Figure 5.**
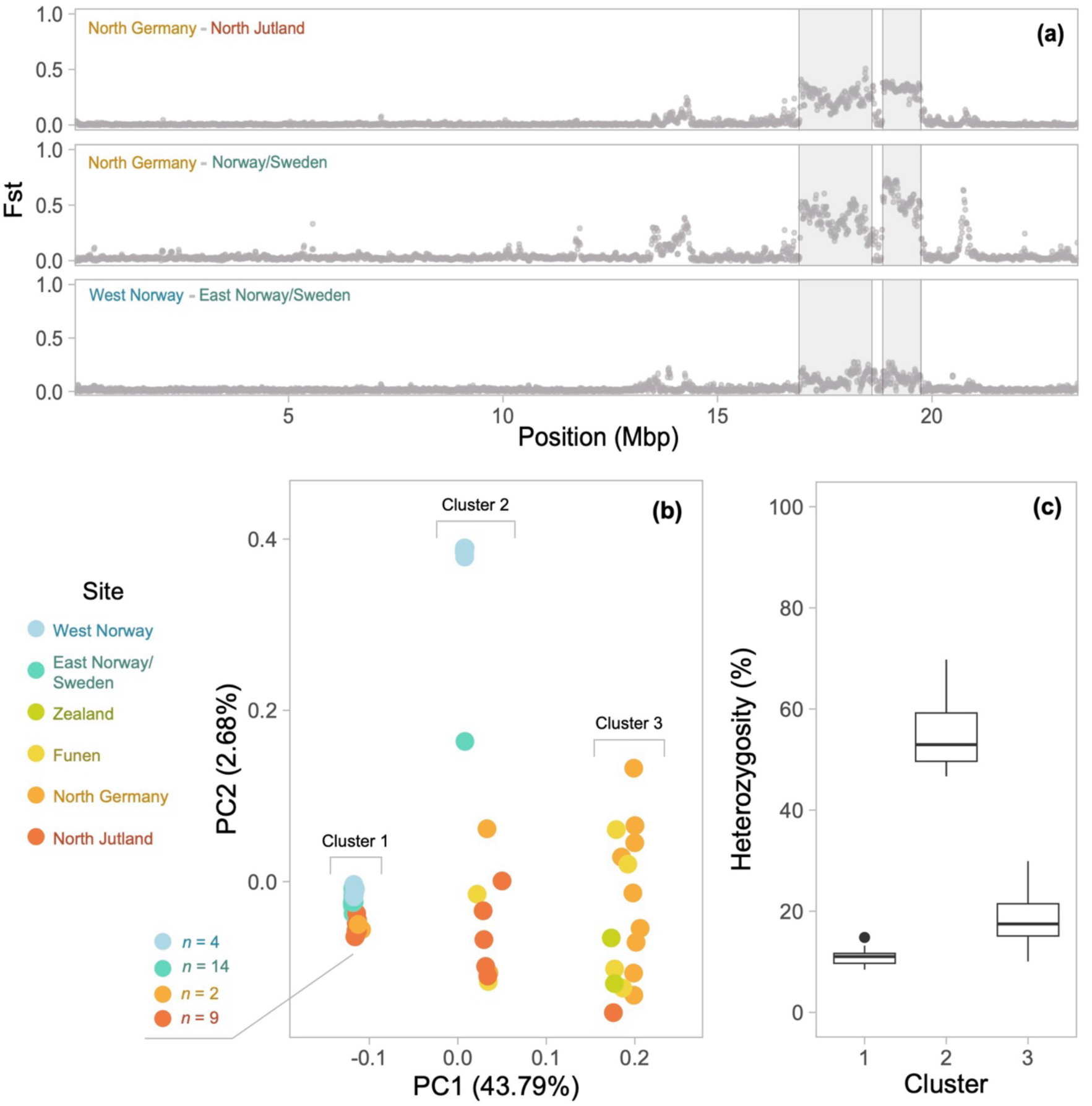
Genomic architecture of the second chromosome of the *B. pascuorum* genome (*n* = 61). Two divergent regions are located between 16.9 - 18.6 Mbp and 18.85 - 19.74 Mbp, and combined contain 23,748 SNPs. **(a)** Pairwise genetic divergence (Fst) between four *B. pascuorum* populations, measured in sliding windows across the second chromosome with a window size 25kb and a step size of 12.5kb. The populations are North Jutland (*n*=15), North Germany (*n*=13), and Norway/Sweden (*n*=25). A pairwise comparison was also conducted within the Norway/Sweden group, comprising Western Norway (*n*=10) and Eastern Norway/Sweden (*n*=15). Coloured labels indicate population pairs. Shaded areas indicate the highly divergent regions from which Principal Component Analysis (PCA) and heterozygosity were measured. **(b)** PCA of the highly divergent regions. Points represent individual samples. Colours indicate sample site: North Germany (*n*=13), North Jutland (*n*=15), Funen (*n*=6), Zealand (*n*=2), Eastern Norway/Sweden (*n*=15) and Western Norway (*n*=10). Percentages indicate the proportion of genomic variation explained by each PC axis. **c)** Boxplot of percentage heterozygosity of the highly divergent regions, compared between the clusters inferred by Figure 5b.

The beginning of chromosome 6 exhibits a narrow region of extremely high divergence in the Germany and Norway/Sweden and the Jutland and Norway/Sweden pairs, with Fst peaking at almost 1 for both pairs. This region occurs between 0.09 and 0.26 Mbp, and contains 85 SNPs. PCA of this region separates all samples into two clusters along PC1, which explains 31% of overall variation, with Norwegian and Swedish samples in one tight cluster and the remaining samples in the other, more divergent cluster. Percentage heterozygosity of this region between sample sites indicates a genotype pattern, with German, North Jutland and Funen samples exhibiting significantly greater heterozygosity than the Norwegian and Swedish samples, as well as the two Zealand samples (W = 542.5, p-value = 0.001) (**Supplementary Figure 13**). PCAs of SNPs in other regions of elevated divergence on chromosome 7 (3-7 Mbp), the beginning of chromosome 13 (0-0.72 Mbp), the end of chromosome 14 (10.3-11.2 Mbp) and on chromosome 16 (2.7-5.8 Mbp) all separated Norwegian and Swedish samples from German and Danish samples along PC1, but local measures of site-specific heterozygosity showed no evidence of distinct biallelic genotypic inheritance (**Supplementary Figures 14-17**).

## Discussion

We uncover distinct population structure in both *B. lapidarius* and *B. pascuorum,* as well as striking contrasts in their genomic architecture. Here, we examine these findings in the biological context of each species, followed by a discussion of their wider consequences for our understanding of bumblebee population structure across fragmented landscapes.

For *B. lapidarius*, population structure was previously detected at a continental scale across Northern Europe between sample sites 1650 km apart, with no evidence of finer substructure (29). Here, we expand on these findings by revealing unprecedented fine-scale population structure throughout our northern European sample range between populations separated by as little as 320km. These patterns are likely a result of combined natural and anthropogenic geographical features impeding dispersal and gene flow between the sites. Divergence of Norwegian and Swedish samples from both Danish and German samples is expected due to the ocean barriers separating the Scandinavian peninsula from continental Europe, which form natural boundaries against bumblebee dispersal and thus drive population structure (18). The divergence between the North Jutland samples and North German samples is less explicable. While the northern Jutlandic island, from which our samples originate, is separated from the rest of continental Denmark by the Limfjord, the most narrow stretches of this boundary are less than 500m wide and well within the estimated *B. lapidarius* queen dispersal capability of at least 5km, and thus could be crossed (50). This high dispersal capability is exemplified by the clustering of the Funen samples, which fall entirely within the North Jutland cluster despite an additional natural barrier represented by the Koldingfjord (a 600m-2km wide channel which divides Funen from continental Denmark). Given that range fragmentation by both urbanisation and agricultural intensification are known drivers of bumblebee population structure (17, 18), more influential dispersal-limiting factor may therefore be the compounding impacts of anthropogenic land-use change across Northern Germany and the Danish peninsula, namely arable land, permanent crops and impervious urban areas (49). Furthermore, we find that population structure in *B. lapidarius* is partly driven by elevated local divergence in narrow genomic regions. These regions occur throughout the genome, and most frequently between the North Germany and Norway/Sweden clusters. One of these small, strongly linked regions exhibits a distinctive three-fold genotype clustering pattern which is indicative of chromosomal rearrangement. *B. lapidarius* does not exhibit any known morphological differentiation across our sample range, and while these elevated peaks suggest local adaptation, their phenotypic consequences remain unknown.

For *B. pascuorum*, overall genomic differentiation between populations is again significant. Population structure in this species has previously been detected at a continental scale, and at a finer scale throughout Sweden consistent with island populations and colour morphology (29, 41). Similarly, we identify fine-scale population structure, yet we also discover substantial spatial variation in structure across our North European range: despite comparable geographical distances, samples from North Jutland and North Germany form distinct genomic clusters, whereas eastern Norwegian and Swedish samples appear much less differentiated from western Norwegian samples. This result is consistent with the contrasting geographical features of the two landscapes which frame the north and south of the Skagerrak and Kattegat straits. While the Norwegian and Swedish mainlands contain numerous water bodies such as fjords, lakes, and rivers, they are not wholly divided by these features. Perhaps more importantly, anthropogenic land-use change occurs at significantly reduced intensities within Norway, much of which remains considerably undisturbed in comparison to the rest of continental Europe (49). This is due in part to the Scandinavian Mountains which dominate a vast proportion of the Norwegian landscape. Indeed, the eastern and western Norwegian sample sites occur on either side of the southernmost part of this mountain range, yet genomic divergence between them remains relatively low. Previous studies have reported similar limited impacts of mountainous landscapes on bumblebee population structure (18, 41).

Furthermore, we here discover that population structure in *B. pascuorum* is in part driven by elevated local divergence in megabase-sized genomic regions. These peaks occur throughout the genome between all three major clusters inferred by the structure analyses, and most strongly and frequently between the North Germany and Norway/Sweden clusters. This local divergence is also present within the Norway/Sweden cluster, specifically, between eastern and western samples. We speculate that some of these genomic regions contribute to the divergence between the four morphological variants which occur throughout the sample range: *B. p. floralis* (North Germany, North Jutland, Funen), *B. p. mniorum* (Zealand) and *B. p. pallidofacies* (East Norway, Sweden) from the *floralis* subgroup, and *B. p. sparreanus* (West Norway) from the *smithianus* subgroup (48). The most explicit of these peaks of divergence, which occupies approximately 2.6 megabases of the second chromosome, also exhibits a distinct three-fold genotype clustering pattern. We hypothesise that this region represents a putative biallelic megabase-scale chromosomal inversion which segregates throughout *B. pascuorum* populations. Large inversions are chromosomal rearrangements which can play pivotal roles in maintaining divergence between ecotypes, underpinning local adaptation, and driving speciation in a range of organisms (51–55). As yet, megabase-sized inversions in bumblebee genomes have solely been recognised at an inter-species level, and overall rates of chromosomal rearrangements over time are thought to be low in comparison to other insect genera (56). This discovery therefore has profound implications for our understanding of intraspecific chromosomal evolution in bumblebees. While the putative inversion is polymorphic in all large sample groups, a clear geographical trend occurs across the three genotypes, with the eastern Norwegian/Swedish samples and North Jutland samples dominated by one homozygous state, the alternate homozygous state exhibited by all but three of North German samples, and the heterozygous state carried by the majority of western Norwegian samples. This spatial pattern, coupled with the highly elevated divergence of the inversion, is indicative of local adaptation.

Several trends are shared between *B. pascuorum* and *B. lapidarius*, such as increased population structure coinciding with oceanic barriers and higher levels of habitat disturbance. While relative genetic divergence (Fst) differs between population pairs, there is very little difference in absolute genetic divergence (Dxy), indicating that population structure in both species is driven by shared polymorphisms which differ in frequency. We also find Norwegian and Swedish populations exhibit significantly lower levels of heterozygosity than those from Denmark and Germany. These patterns are indicative of post-glacial colonisation of the Fennoscandian peninsula, suggesting that this divergence is therefore of fairly recent origin.

Nonetheless, we also identify distinct genomic contrasts between the two species: *B. pascuorum* exhibits stronger structure, as well as significantly higher indicators of recent inbreeding. Although .similar species-specific patterns have been observed in bumblebees sampled from Ireland and Great Britain, with significant structure identified in the montane *Bombus monticola*, and little structure occurring in the widespread *Bombus pratorum* (40), these results are unsurprising due to their contrasting ecologies. We also find larger regions of elevated local divergence in *B. pascuorum*, and while *B. lapidarius* also exhibits several regions with elevated divergence, including one with signs of biallelic inheritance, these regions are relatively small, and occur far less frequently throughout the genome. It is unclear which biological factors underlie these differences. It is possible that the greater structure in *B. pascuorum* may be evidence of earlier range expansion into Northern Europe and the Scandinavian peninsula following the Last Glacial Maximum compared to *B. lapidarius*. Alternatively, such contrasting patterns of genomic structure may be caused by differential sensitivities to anthropogenic land-use change, and associated stressors such as pollution, perhaps due to adaptive differences in foraging or nesting ecology. Indeed, *B. lapidarius* has been known to inhabit both semi-urban and agricultural landscapes, and therefore appears to be among the most resilient bumblebees to anthropogenic pressures (45, 46). The flight distances of *B. lapidarius* queens have also been estimated to be greater than those of *B. pascuorum*, suggesting a comparatively higher dispersal capability (50). The strong population structure observed between *B. lapidarius* populations in Denmark and Germany is therefore evidence for significant dispersal barriers throughout this anthropogenic landscape.

## Conclusions

Here, we use whole genome data to present definitive evidence of fine-scale population structure in two widely distributed bumblebee species (*B. pascuorum* and *B. lapidarius*) throughout northern Europe. Several conclusions can be drawn from our results. First, these levels of structure are remarkable given the relatively recent colonisation of this region following the Last Glacial Maximum, as well as the wide distributions, high dispersal capacity and relatively high abundance of both study species. Importantly, our observations imply that significantly more population structure can be expected throughout the broad European ranges of these species, which are fragmented by similar or higher levels of anthropogenic land-use as encountered in our sample range. It is likely that many regions harbour local bumblebee populations that are not connected by high levels gene flow (40), and even greater population structure can be expected to occur in rare or threatened species, such as *Bombus muscorum* (24) or *Bombus sylvarum* (35). Such populations are sensitive to the impacts of habitat loss and range fragmentation, and may therefore be at increased risk of local extinction. Continued assessments of the range-wide genomic diversity of these species are therefore crucial for the effective identification of target populations for conservation management. Secondly, while we observe overlapping patterns between the population structure of *B. pascuorum* and *B. lapidarius*, such as those associated with natural oceanic barriers to gene flow, there are also several trends through which the two species significantly differ. These differences illustrate that population genomic structure in bumblebees is species-specific, even for species with similar distributions, abundance and spread. Thus, bumblebee population structure is not entirely predictable from one species to the next. There is therefore an urgent need to apply population genomic studies to a greater range of species throughout the bumblebee genus in order to bring our understanding of bumblebee connectivity to a greater taxonomic resolution. Finally, by analysing millions of genome-wide polymorphisms, we discover specific regions of highly elevated genomic structure that are indicative of local adaptation, shedding light on bumblebee subpopulation diversity and adaptive potential. Altogether, our results are of fundamental importance for our overall understanding of bumblebee genomic variation, population connectivity and conservation.

## Materials & Methods

### Sampling, DNA extraction and whole genome sequencing

*B. pascuorum* individuals (*n*=73) and *B. lapidarius* individuals (*n*=55) were sampled between 2016-2022 from up to 7 sites spread between the southern Scandinavian peninsula and Germany: North Germany, North Jutland, Funen, Zealand, the Swedish coast, East Norway and West Norway. DNA was extracted from 1 or 2 bumblebee legs using the DNeasy Blood & Tissue kit (Qiagen) following the manufacturer’s protocol. Genomic libraries were built using an in-house developed tagmentation-based library preparation method (57), and then sequenced by the Norwegian Sequencing Center. Initially, 44 *B. pascuorum* samples and 52 *B. lapidarius* samples were sequenced using an Illumina Novaseq 6000 SP flowcell, generating 923,253,536 read pairs passing filters. All samples were then sequenced using an Illumina 6000 S4 flowcell and generated 4,680,209,690 read pairs passing filters. For more information about DNA extraction, library preparation and sequencing, see the **Supplementary Text**.

### Variant calling, data cleaning and quality control

Base calling and demultiplexing for all samples were performed using RTA v3.4.4 and bcl2fastq v2.20.0.422. Following the removal of sequencing adaptors with AdapterRemoval/2.3.1 (58), read data were aligned to the draft assembly of BomPasc1.1 (59) and the contig-level assembly of BomLapEIv1 (60) using PALEOMIX v1.2.14 (61) and the BWA/0.7.17 mem algorithm (62). All reads with a minimum MapQ value of 15 were retained and considered endogenous. Clonal reads were removed using Picard tools v.3.1.1 (63). Variants were called from the bam files with BCFtools v1.15.1 using the *mpileup* and multiallelic *call* tools. VCFs were then filtered using VCFtools v0.1.16, excluding indels, sites with over 20% missing data, and sites with a Quality score lower than 20. Only sites with a mean depth of between 5X and 40X across all individuals were retained, as well as all genotypes with a depth greater than 5X. A minimum allele frequency threshold of 0.05 was also applied. All individuals with a mean coverage lower than 5X (4 *B. pascuorum* and one *B. lapidarius*) and mean missingness of over 25% (3 *B. pascuorum* and 3 *B. lapidarius*) were removed. Two pairs of individuals in the *B. pascuorum* dataset and one pair in the *B. lapidarius* dataset were found to be highly related (KING kinship coefficient < 0.2; (64)), and in each instance the individual with lower mean coverage was removed. Seven mistakenly sampled individuals consisting of three males and four *Bombus sylvarum* bumblebees were also removed from the dataset. In addition, one *B. lapidarius* sample exhibited unusually high heterozygosity (F=-0.4), likely due to contamination, and was thus removed.

Following data cleaning, filtering and quality control, the final dataset retained for all population genomic analyses consisted of 2,839,545 SNPs for *B. pascuorum*, with 61 individuals from 7 sites: North Germany (*n*=13), North Jutland (*n*=15), Funen (*n*=6), Zealand (*n*=2), the Swedish coast (*n*=4), East Norway (*n*=11) and West Norway (*n*=10) (**Figure 1**). The final *B. lapidarius* dataset consisted of 1,464,934 SNPs, with 45 individuals from 6 sites: North Germany (*n*=11), North Jutland (*n*=13), Funen (*n*=3), Zealand (*n*=2), the Swedish coast (*n*=2) and East Norway (*n*=15) (**Figure 1**).

### Clustering and structure analyses

Prior to all clustering and structure analyses, the dataset was pruned for linkage disequilibrium using PLINK v2.0, excluding all sites with an r^2^ value greater than 0.3. 1,863,457 *B. pascuorum* SNPs and 698,786 *B. lapidarius* SNPs were retained following linkage pruning. Principal component analysis (PCA) was conducted using the *smartPCA* algorithm as implemented in EIGENSOFT v45.2.1. Population structure analysis was conducted with ADMIXTURE v1.3.0, which was run 10 times for K=1 to K=5 with a different random seed each time. For each value of K, the ancestry fraction estimates with the highest log-likelihood were selected from the 10 outputs and evaluated using evalAdmix v0.95. Between-group statistical comparisons of model fit for each K value were conducted using Kruskal-Wallis and pairwise Wilcoxon tests.

### Genomic diversity and inbreeding

Hudson’s pairwise Fst between populations indicated by prior clustering analyses were carried out in PLINK v2.0. Pairwise Dxy between populations were carried out using the popgenWindows python script by S. Martin (65). *B. lapidarius* populations were grouped as follows: Germany (*n*=10), North Jutland/Funen (*n*=16), and Norway/Sweden (*n*=17). Most *B. pascuorum* sample sites formed separate clusters and were retained as groups, except for Norwegian and Swedish individuals which formed one cluster and were therefore grouped together. Zealand individuals did not group with any of the major clusters in either species, and were thus excluded from this analysis due to their small sample sizes (*n*=2). To generate Tajima’s D, no minimum allele frequency threshold was used, resulting in 10,544,756 SNPs for *B. lapidarius*, and 18,034,694 SNPs for *B. pascuorum.* Tajima’s D was calculated in 50,000bp bins for all clusters except Zealand using VCFtools v0.1.16. F statistics were generated using VCFtools v0.1.16. Genome-wide heterozygosity was measured using BCFtools v1.15.1 (for detailed method, see **Supplementary Text**). Runs of homozygosity (RoH) were identified using PLINK v1.90. RoHs were first estimated from the unpruned dataset, followed by data lightly pruned for linkage disequilibrium (r2 > 0.9), also using PLINK, as recommended by (66) for improved detection of autozygous RoHs. RoH results generated from both pruned and unpruned data did not differ, and the unpruned dataset was therefore retained. As per (41), a minimum RoH length threshold of 10kb was applied, containing at least 50 SNPs with a minimum density of 1 SNP per 50kb. A maximum of 3 missing sites and 0 heterozygotes were allowed per 50kb sliding window, and a maximum gap of 1000 kb between consecutive SNPs was required for them to be considered separate RoHs.

### Genome-wide divergence and genomic architecture

Pairwise Fst values between populations inferred by prior clustering analyses were calculated in sliding windows across the genome using VCFTools v0.1.16, with a window size of 25kb and a step size of 12.5kb. Given that the reference genome used for *B. lapidarius* is currently contig-level with no assembled chromosomes, the first 95 unplaced scaffolds were used, comprising 1,390,497 SNPs and occupying 72% of the full genome. *B. lapidarius* populations were defined as North Jutland/Funen individuals (*n*=16), German individuals (*n*=10) and Norwegian/Swedish individuals (*n=*17). The two Zealand individuals, which did not group with any major cluster, were excluded from the analysis due to their small sample size (*n*=2). For *B. pascuorum,* all unplaced scaffolds were excluded from the analysis, and solely the 17 assembled chromosomes were retained, comprising 2,784,282 SNPs. *B. pascuorum* populations were defined as North Jutland individuals (*n*=15), German individuals (*n*=13) and Norwegian/Swedish individuals (*n*=25). Individuals from Funen and Zealand did not group with any major cluster, were excluded from the analysis due to their small sample size (*n*=6; *n*=2). PCAs of highly variable regions were conducted in PLINK v2.0 without pruning for linkage disequilibrium, and measures of heterozygosity were conducted in VCFtools v0.1.16. All sample sites were included in PCAs and measures of heterozygosity. Fst values are considered outliers when they exceed the 99^th^ percentile of Fst frequency distribution (assuming normal distribution).

### Statistical analyses

Tracy-Widom tests for PC axes were conducted using *smartPCA*. Between-group statistical comparisons of model fit for each K value were conducted using Kruskal-Wallis and pairwise Wilcoxon tests. Between-site comparisons of individual F statistics, mean heterozygosity, RoH length and percentage of the genome covered by RoHs were performed using analysis of variance tests and post-hoc Tukey Honest Significant Difference tests for data meeting assumptions required for parametric tests (normal distribution and homogeneity of variance, which was tested using Bartlett’s tests). Pairwise parametric comparisons of these measures between the two species were conducted using t-tests. Data which failed to meet parametric test assumptions were analysed using Kruskal-Wallis tests and post-hoc Dunn tests for multiple-group comparisons, and pairwise Wilcoxon tests for pairwise comparisons. All statistical analyses were carried out in base R v4.3.2.

## Data Availability Statement

All raw sequence read data are available under PRJEB65580 and PRJEB74745 at the European Nucleotide Archive (ENA, https://www.ebi.ac.uk; **Supplementary Table 2**).

## Supplementary Figures

**Supplementary Figure 1.**
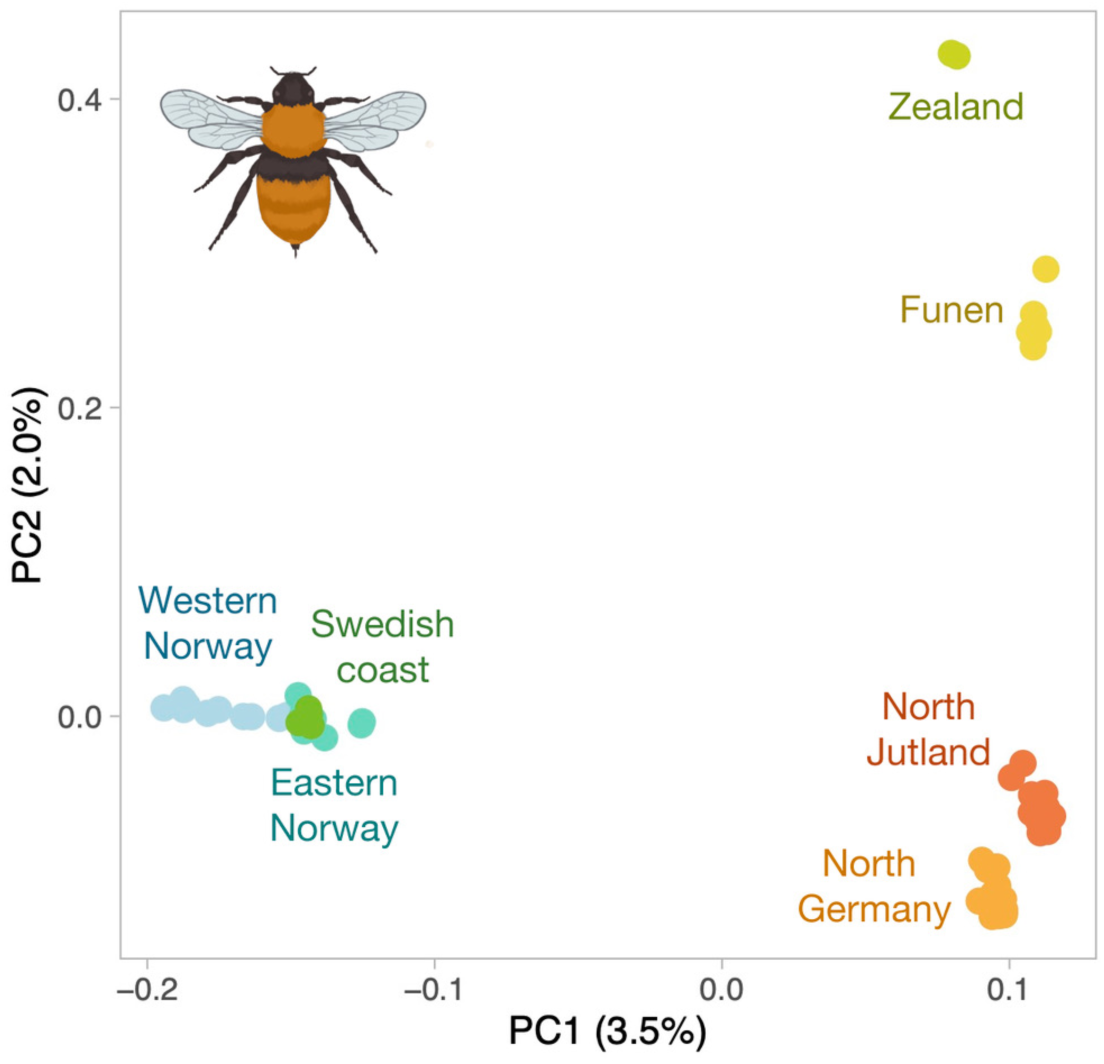
Distinct genome-wide population structure in *Bombus pascuorum* across northern Europe. Principal component analysis (PCA) plot of 1,863,457 SNPs from 61 *B. pascuorum* individuals sampled across southern Norway, southern Sweden, Denmark and north Germany. PC1 and PC3 are presented. Points represent individual samples and colours indicate sample site assignments. Percentages indicate the proportion of genomic variation explained by each PC axis.

**Supplementary Figure 2.**
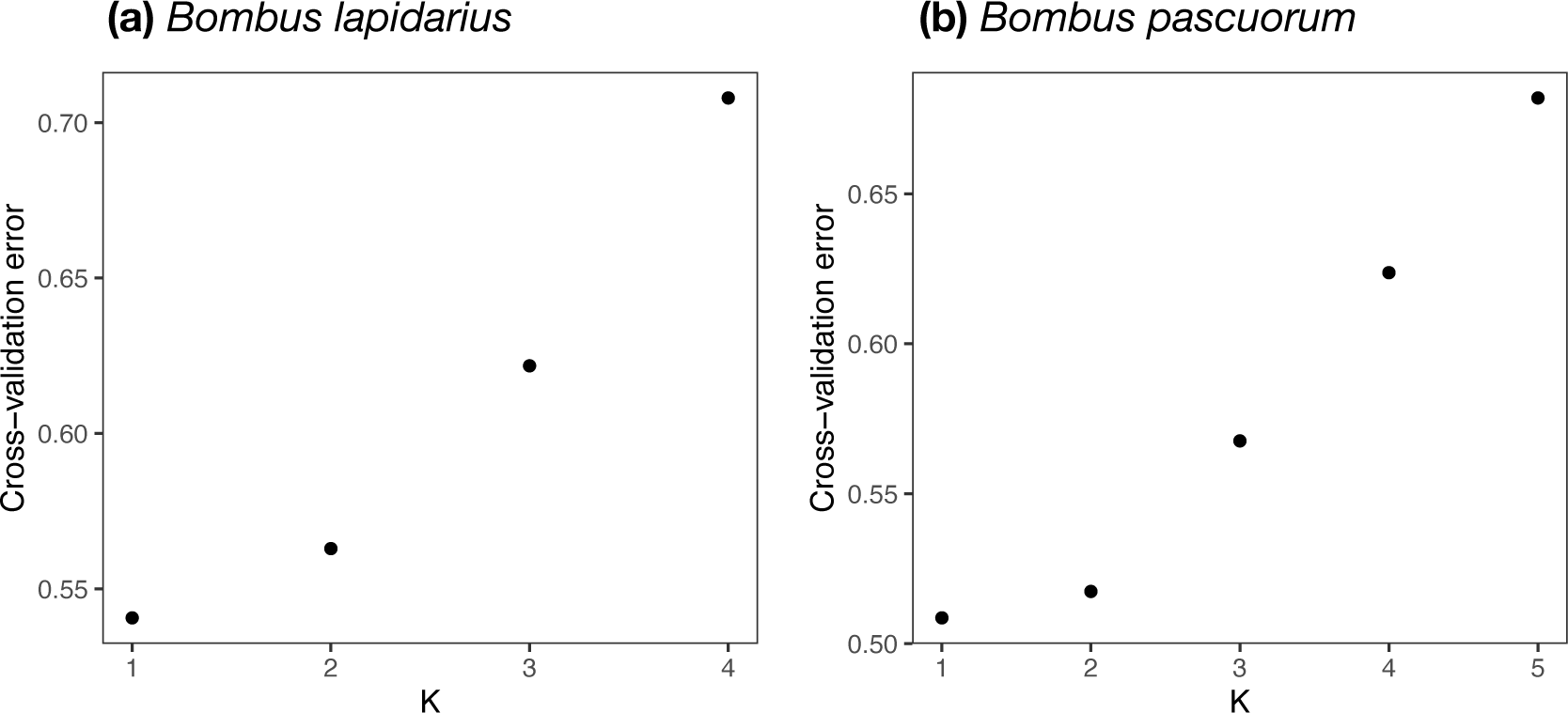
Cross-validation plots for ADMIXTURE analysis of (a) 45 *Bombus lapidarius* individuals and (b) 61 *Bombus pascuorum* individuals sampled across southern Norway, southern Sweden, Denmark and north Germany. **(a)** Ancestry proportions were estimated from ADMIXTURE analysis of 698,786 SNPs for K=2 - K=4. **(b)** Ancestry proportions were estimated from ADMIXTURE analysis of 1,863,457 SNPs for K=2 - K=5.

**Supplementary Figure 3.**
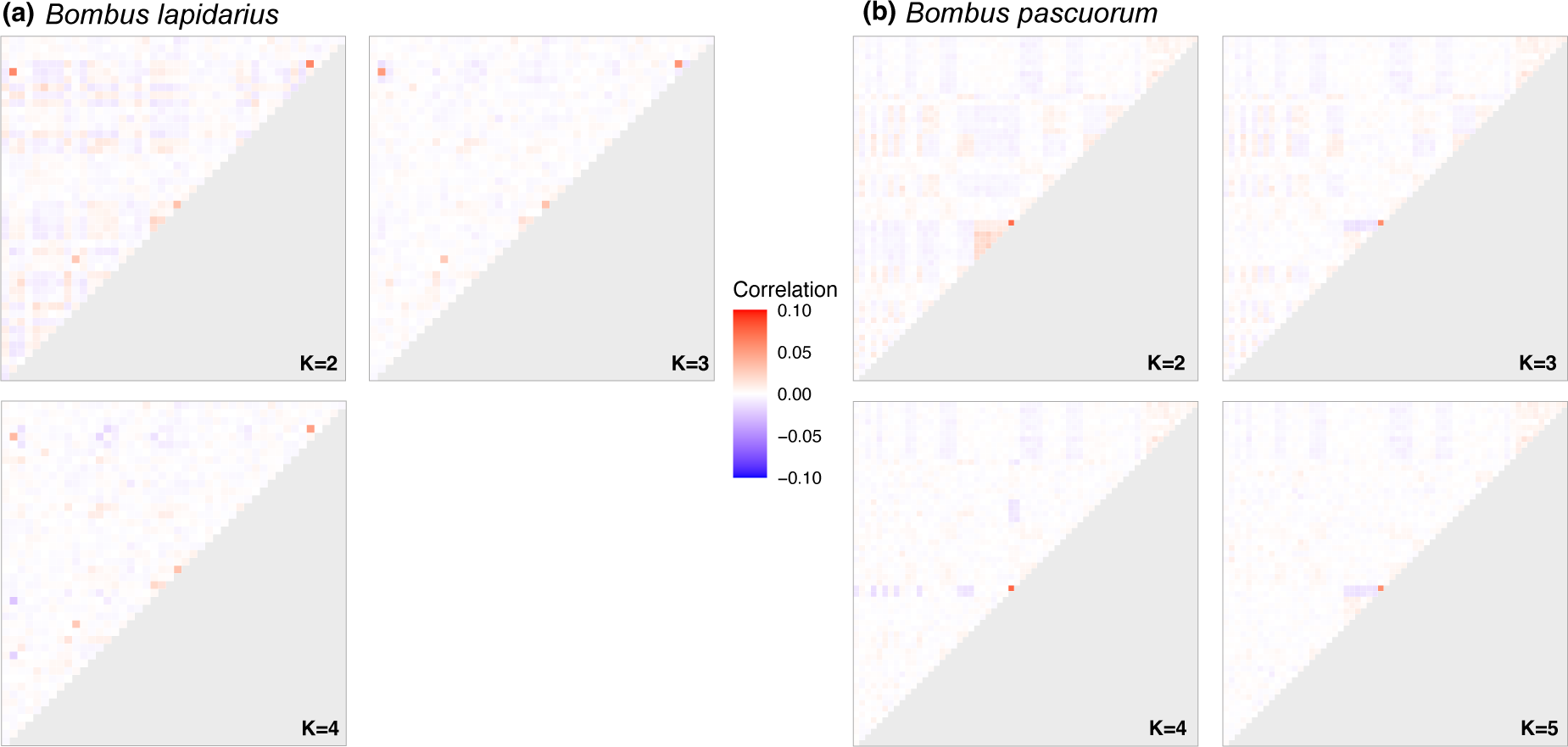
Correlation matrix plots generated by evalAdmix model validation for ADMIXTURE analysis of (a) 45 *Bombus lapidarius* individuals and (b) 61 *Bombus pascuorum* individuals sampled across southern Norway, southern Sweden, Denmark and north Germany. **(a)** Model validation of ancestry proportions estimated from ADMIXTURE analysis of 698,786 SNPs for K=2 - K=4. **(b)** Model validation of ancestry proportions estimated from ADMIXTURE analysis of 1,863,457 SNPs for K=2 - K=5. Each column and row represents a sample, arranged in the same order on each axis. Correlation coefficients fall between -0.1 (red) and 0.1 (blue), with white representing a correlation coefficient of 0 (no correlation).

**Supplementary Figure 4.**
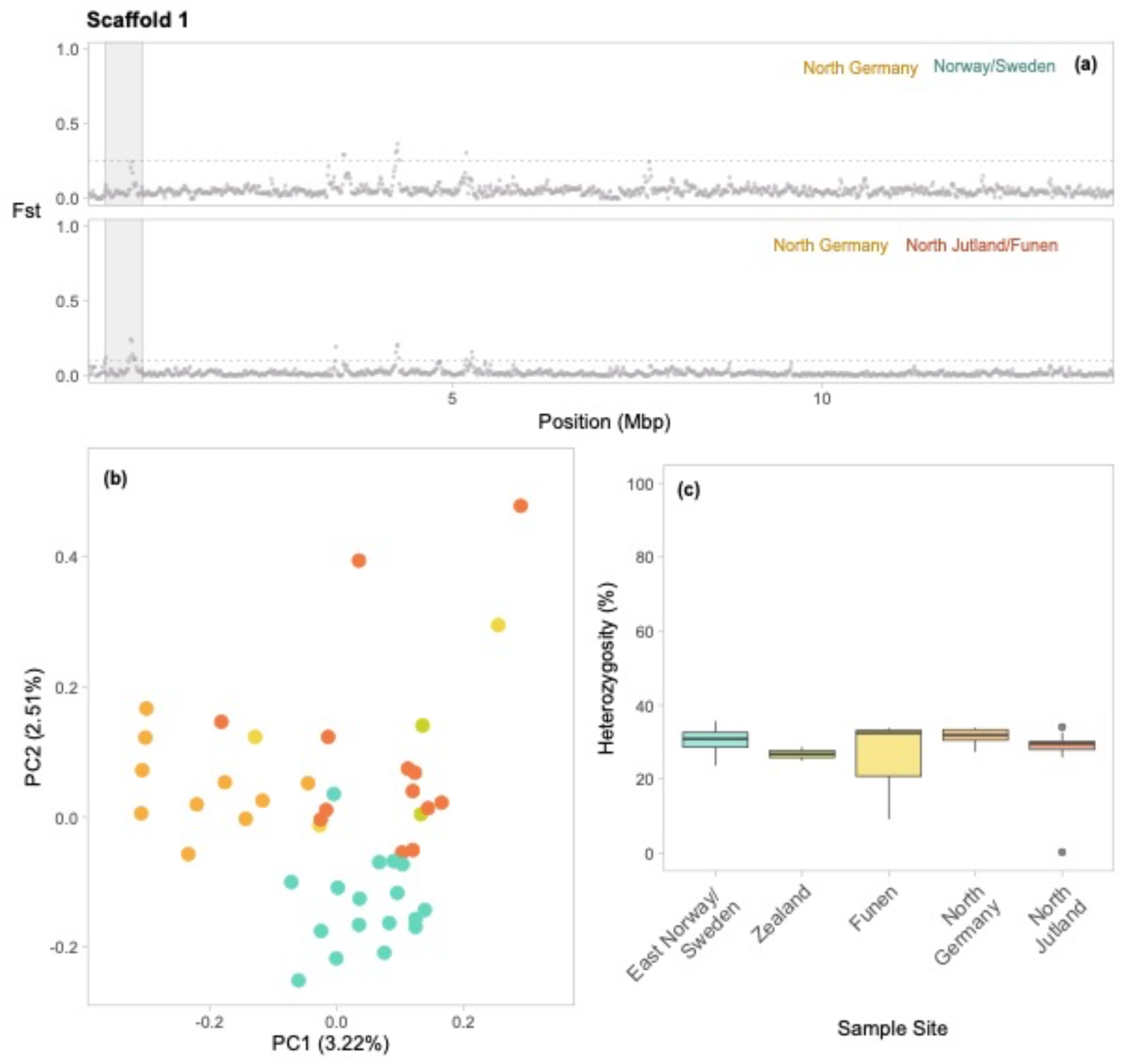
Genomic architecture of the first scaffold of the *B. lapidarius* genome (*n* = 45). The highly divergent region is located between 0.3 and 0.8 Mbp, and contains 3393 SNPs. **(a)** Pairwise genetic divergence (Fst) between three *B. lapidarius* populations, measured in sliding windows across the second chromosome with a window size 25kb and a step size of 12.5kb. The populations are North Jutland/Funen (*n*=16), North Germany (*n*=11), and East Norway/Sweden (*n*=17). Coloured labels indicate population pairs. Shaded areas indicate the highly divergent region from which Principal Component Analysis (PCA) and heterozygosity were measured. **(b)** PCA of the highly divergent region. Points represent individual samples. Colours indicate sample site: North Germany (*n*=11), North Jutland (*n*=13), Funen (*n*=3), Zealand (*n*=2), and East Norway/Sweden (*n*=17). Percentages indicate the proportion of genomic variation explained by each PC axis. **(c)** Boxplot of percentage heterozygosity of the highly divergent regions, compared between the sample sites.

**Supplementary Figure 5.**
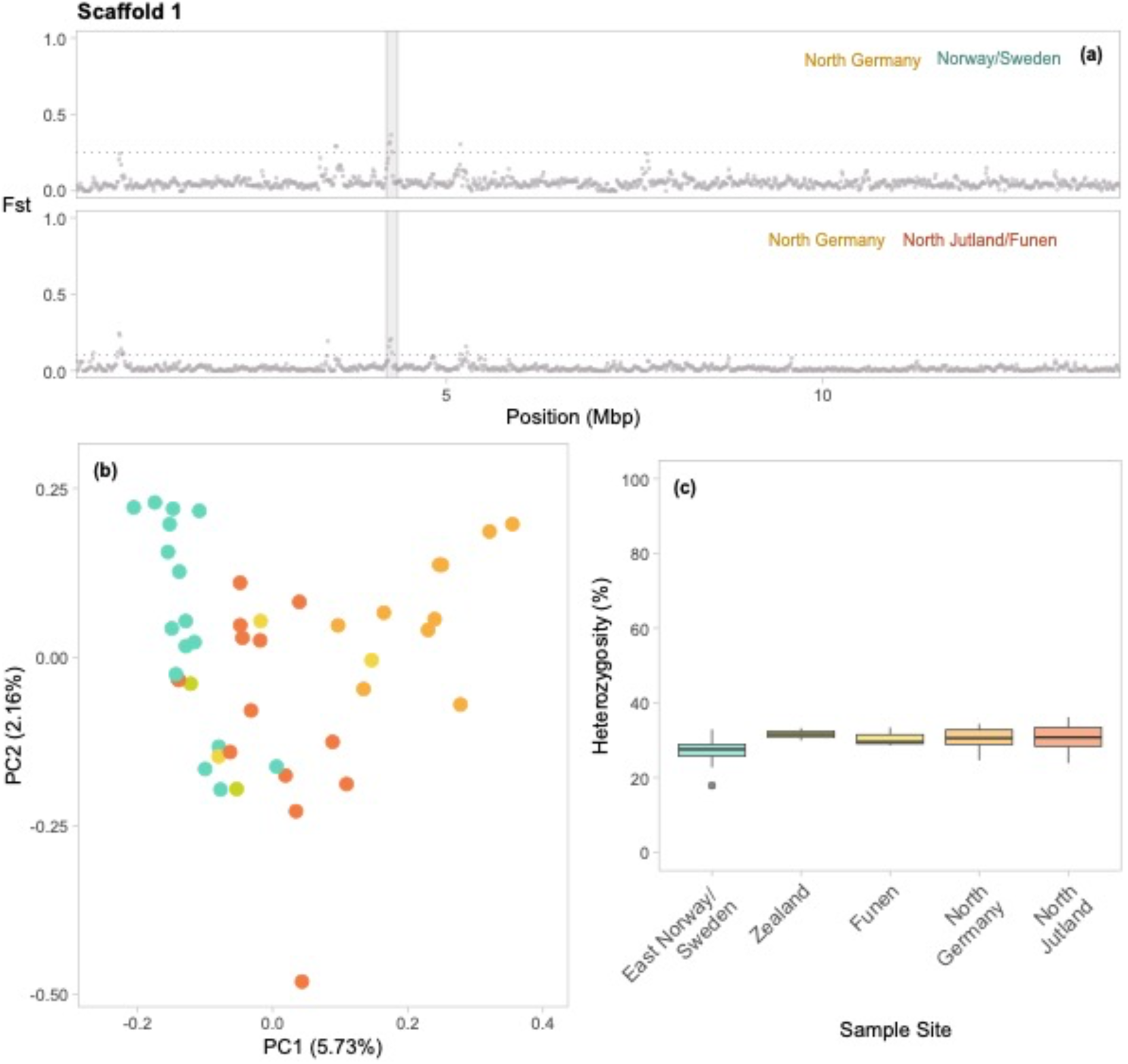
Genomic architecture of the first scaffold of the *B. lapidarius* genome (*n* = 45). The highly divergent region is located between 4.20 and 4.35 Mbp, and contains 994 SNPs. **(a)** Pairwise genetic divergence (Fst) between three *B. lapidarius* populations, measured in sliding windows across the second chromosome with a window size 25kb and a step size of 12.5kb. The populations are North Jutland/Funen (*n*=16), North Germany (*n*=11), and East Norway/Sweden (*n*=17). Coloured labels indicate population pairs. Dotted lines represent the 99^th^ percentile of each Fst frequency distribution. Shaded areas indicate the highly divergent region from which Principal Component Analysis (PCA) and heterozygosity were measured. **(b)** PCA of the highly divergent region. Points represent individual samples. Colours indicate sample site: North Germany (*n*=11), North Jutland (*n*=13), Funen (*n*=3), Zealand (*n*=2), and East Norway/Sweden (*n*=17). Percentages indicate the proportion of genomic variation explained by each PC axis. **(c)** Boxplot of percentage heterozygosity of the highly divergent regions, compared between the sample sites.

**Supplementary Figure 6.**
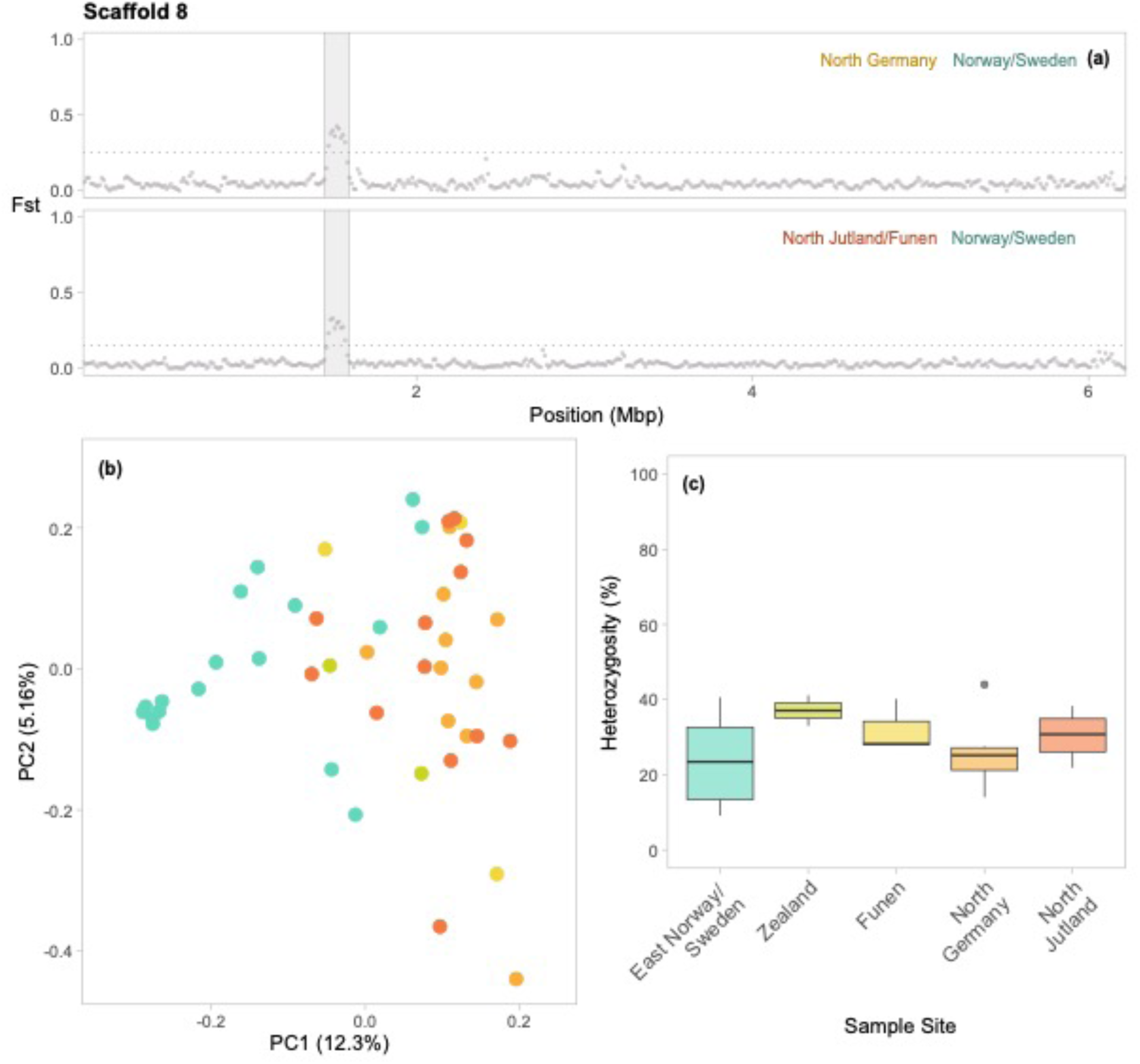
Genomic architecture of the 8th scaffold of the *B. lapidarius* genome (*n* = 45). The highly divergent region is located between 1.45 and 1.60 Mbp, and contains 209 SNPs. **(a)** Pairwise genetic divergence (Fst) between three *B. lapidarius* populations, measured in sliding windows across the second chromosome with a window size 25kb and a step size of 12.5kb. The populations are North Jutland/Funen (*n*=16), North Germany (*n*=11), and East Norway/Sweden (*n*=17). Coloured labels indicate population pairs. Dotted lines represent the 99^th^ percentile of each Fst frequency distribution. Shaded areas indicate the highly divergent region from which Principal Component Analysis (PCA) and heterozygosity were measured. **(b)** PCA of the highly divergent region. Points represent individual samples. Colours indicate sample site: North Germany (*n*=11), North Jutland (*n*=13), Funen (*n*=3), Zealand (*n*=2), and East Norway/Sweden (*n*=17). Percentages indicate the proportion of genomic variation explained by each PC axis. **(c)** Boxplot of percentage heterozygosity of the highly divergent regions, compared between the sample sites.

**Supplementary Figure 7.**
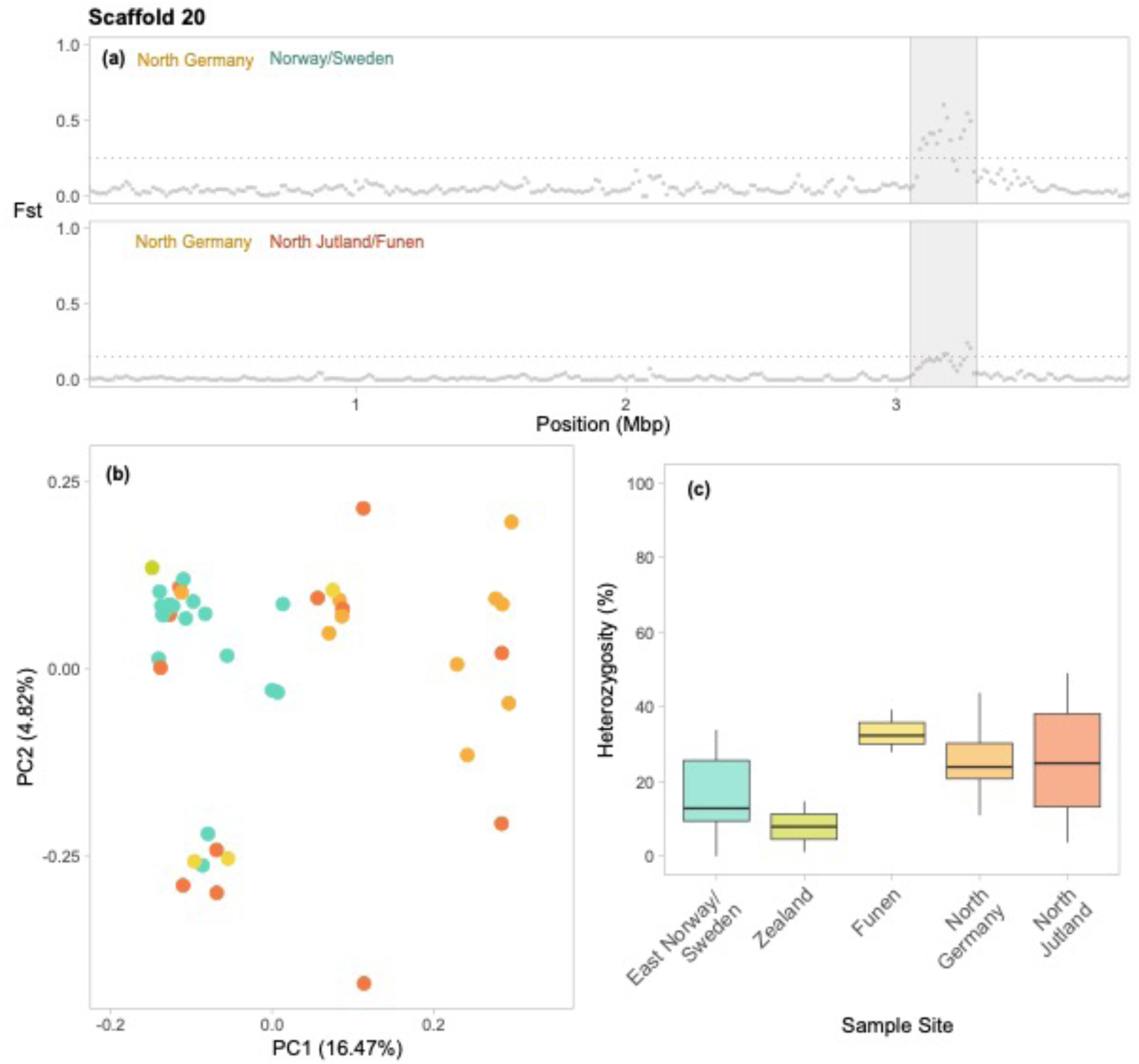
Genomic architecture of the 20th scaffold of the *B. lapidarius* genome (*n* = 45). The highly divergent region is located between 3.05 and 3.30 Mbp, and contains 265 SNPs. **(a)** Pairwise genetic divergence (Fst) between three *B. lapidarius* populations, measured in sliding windows across the second chromosome with a window size 25kb and a step size of 12.5kb. The populations are North Jutland/Funen (*n*=16), North Germany (*n*=11), and East Norway/Sweden (*n*=17). Coloured labels indicate population pairs. Dotted lines represent the 99^th^ percentile of each Fst frequency distribution. Shaded areas indicate the highly divergent region from which Principal Component Analysis (PCA) and heterozygosity were measured. **(b)** PCA of the highly divergent region. Points represent individual samples. Colours indicate sample site: North Germany (*n*=11), North Jutland (*n*=13), Funen (*n*=3), Zealand (*n*=2), and East Norway/Sweden (*n*=17). Percentages indicate the proportion of genomic variation explained by each PC axis. **(c)** Boxplot of percentage heterozygosity of the highly divergent regions, compared between the sample sites.

**Supplementary Figure 8.**
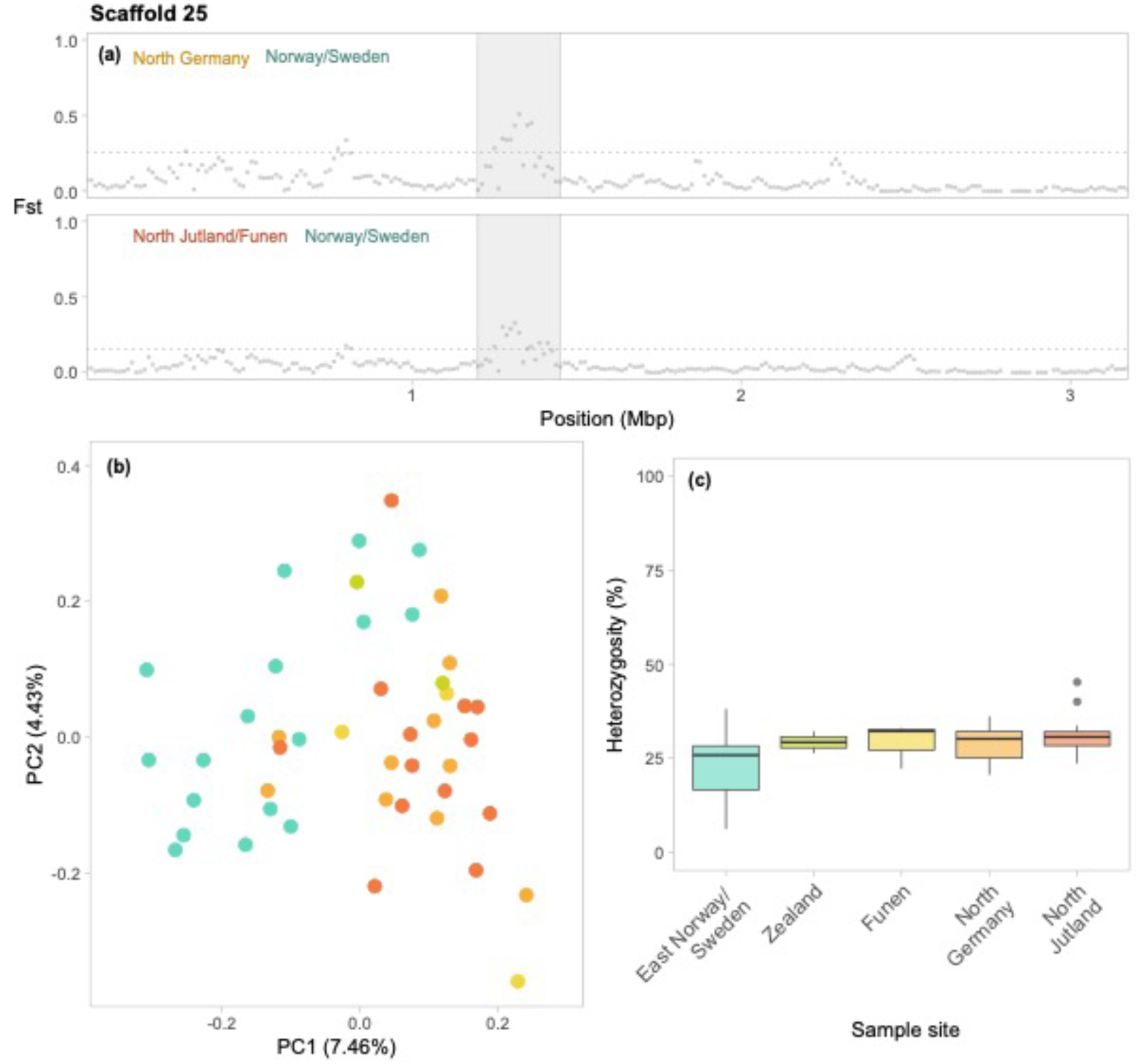
Genomic architecture of the 25th scaffold of the *B. lapidarius* genome (*n* = 45). The highly divergent region is located between 1.20 and 1.45 Mbp, and contains 267 SNPs. **(a)** Pairwise genetic divergence (Fst) between three *B. lapidarius* populations, measured in sliding windows across the second chromosome with a window size 25kb and a step size of 12.5kb. The populations are North Jutland/Funen (*n*=16), North Germany (*n*=11), and East Norway/Sweden (*n*=17). Coloured labels indicate population pairs. Dotted lines represent the 99^th^ percentile of each Fst frequency distribution. Shaded areas indicate the highly divergent region from which Principal Component Analysis (PCA) and heterozygosity were measured. **(b)** PCA of the highly divergent region. Points represent individual samples. Colours indicate sample site: North Germany (*n*=11), North Jutland (*n*=13), Funen (*n*=3), Zealand (*n*=2), and East Norway/Sweden (*n*=17). Percentages indicate the proportion of genomic variation explained by each PC axis. **(c)** Boxplot of percentage heterozygosity of the highly divergent regions, compared between the sample sites.

**Supplementary Figure 9.**
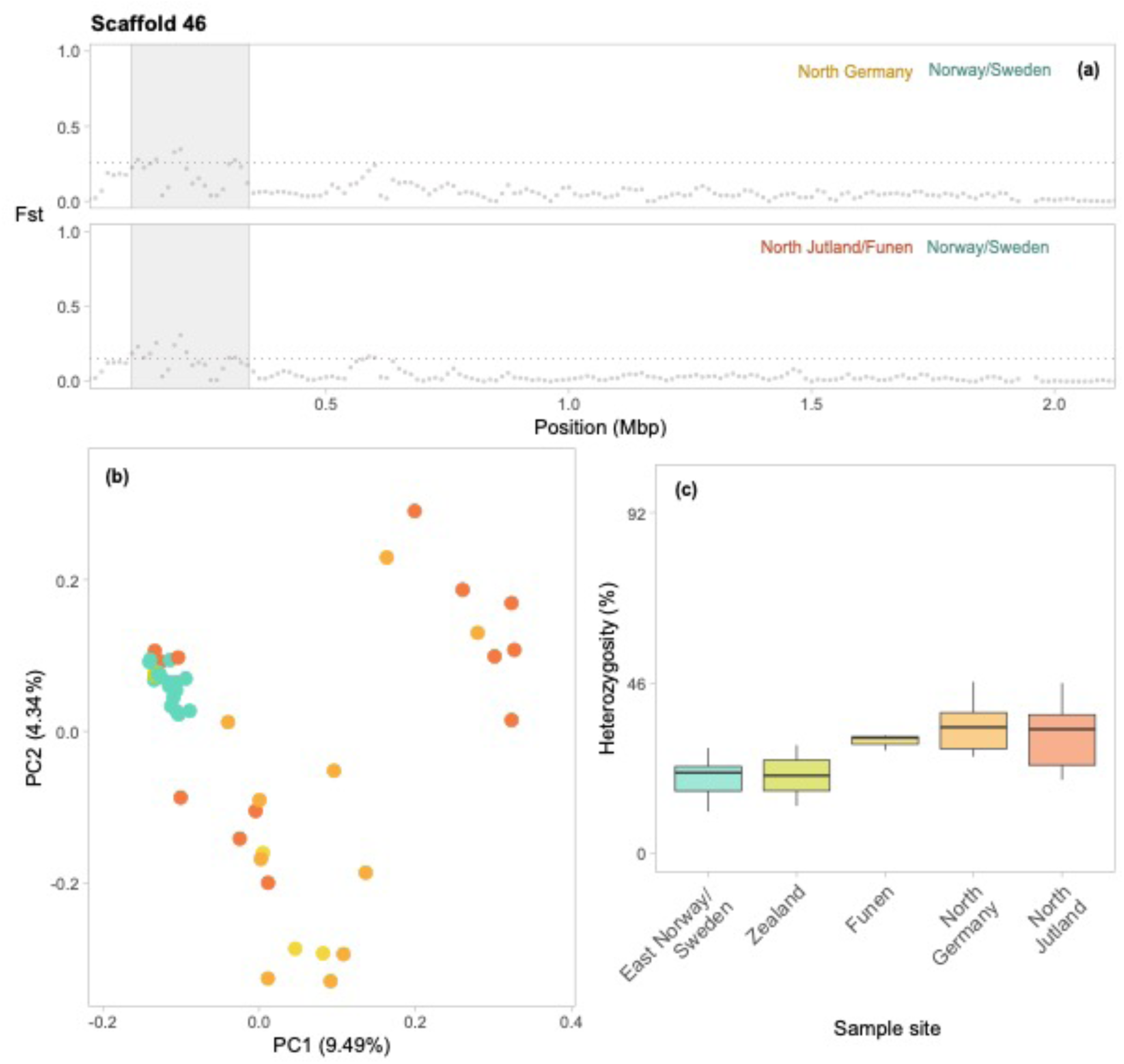
Genomic architecture of the 46th scaffold of the *B. lapidarius* genome (*n* = 45). The highly divergent region is located between 0.10 and 0.34 Mbp, and contains 256 SNPs. **(a)** Pairwise genetic divergence (Fst) between three *B. lapidarius* populations, measured in sliding windows across the second chromosome with a window size 25kb and a step size of 12.5kb. The populations are North Jutland/Funen (*n*=16), North Germany (*n*=11), and East Norway/Sweden (*n*=17). Coloured labels indicate population pairs. Dotted lines represent the 99^th^ percentile of each Fst frequency distribution. Shaded areas indicate the highly divergent region from which Principal Component Analysis (PCA) and heterozygosity were measured. **(b)** PCA of the highly divergent region. Points represent individual samples. Colours indicate sample site: North Germany (*n*=11), North Jutland (*n*=13), Funen (*n*=3), Zealand (*n*=2), and East Norway/Sweden (*n*=17). Percentages indicate the proportion of genomic variation explained by each PC axis. **(c)** Boxplot of percentage heterozygosity of the highly divergent regions, compared between the sample sites.

**Supplementary Figure 10.**
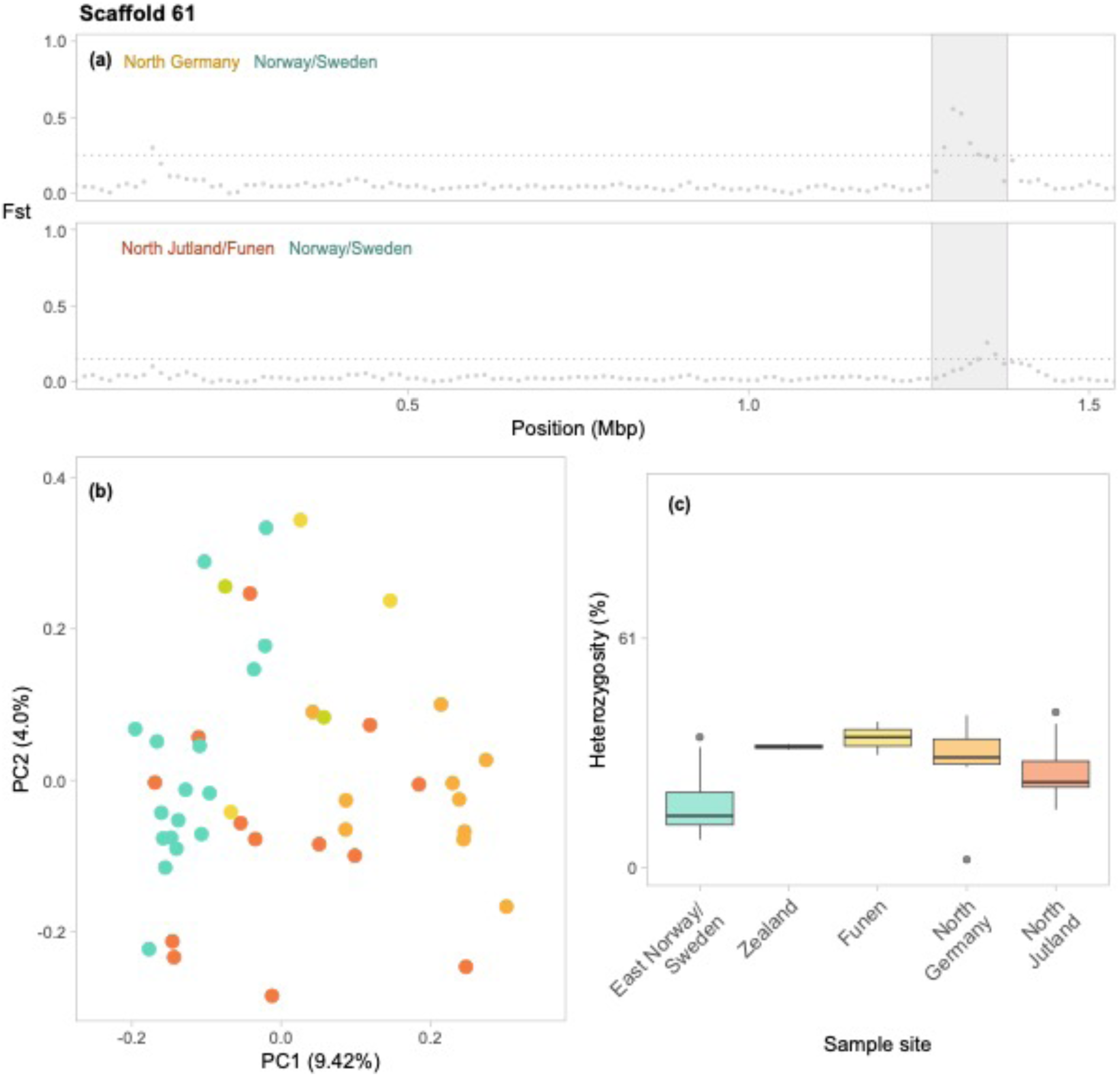
Genomic architecture of the 61st scaffold of the *B. lapidarius* genome (*n* = 45). The highly divergent region is located between 1.27 and 1.38 Mbp, and contains 150 SNPs. **(a)** Pairwise genetic divergence (Fst) between three *B. lapidarius* populations, measured in sliding windows across the second chromosome with a window size 25kb and a step size of 12.5kb. The populations are North Jutland/Funen (*n*=16), North Germany (*n*=11), and East Norway/Sweden (*n*=17). Coloured labels indicate population pairs. Dotted lines represent the 99^th^ percentile of each Fst frequency distribution. Shaded areas indicate the highly divergent region from which Principal Component Analysis (PCA) and heterozygosity were measured. **(b)** PCA of the highly divergent region. Points represent individual samples. Colours indicate sample site: North Germany (*n*=11), North Jutland (*n*=13), Funen (*n*=3), Zealand (*n*=2), and East Norway/Sweden (*n*=17). Percentages indicate the proportion of genomic variation explained by each PC axis. **(c)** Boxplot of percentage heterozygosity of the highly divergent regions, compared between the sample sites.

**Supplementary Figure 11.**
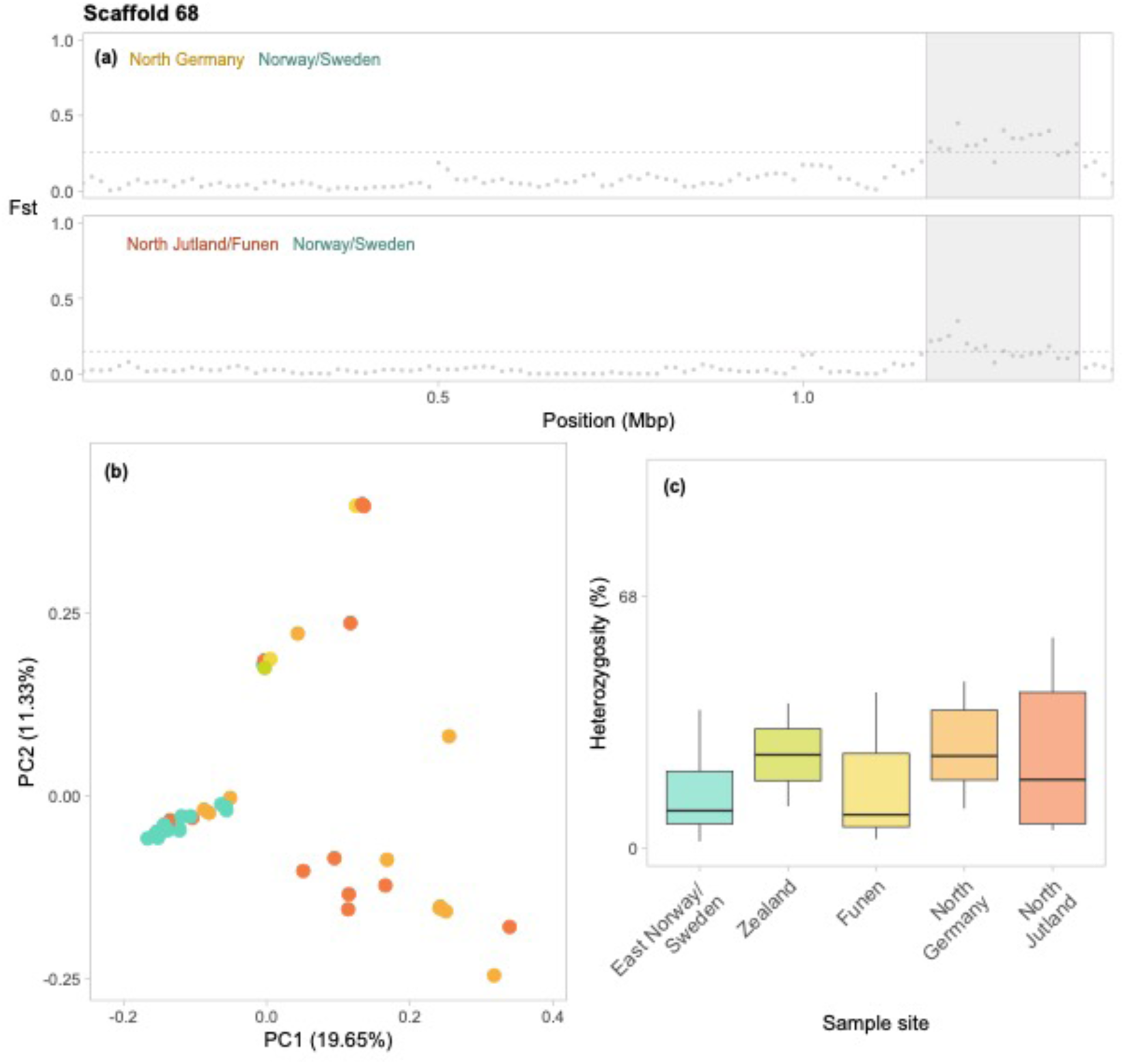
Genomic architecture of the 68th scaffold of the *B. lapidarius* genome (*n* = 45). The highly divergent region is located between 1.17 and 1.38 Mbp, and contains 169 SNPs. **(a)** Pairwise genetic divergence (Fst) between three *B. lapidarius* populations, measured in sliding windows across the second chromosome with a window size 25kb and a step size of 12.5kb. The populations are North Jutland/Funen (*n*=16), North Germany (*n*=11), and East Norway/Sweden (*n*=17). Coloured labels indicate population pairs. Dotted lines represent the 99^th^ percentile of each Fst frequency distribution. Shaded areas indicate the highly divergent region from which Principal Component Analysis (PCA) and heterozygosity were measured. **(b)** PCA of the highly divergent region. Points represent individual samples. Colours indicate sample site: North Germany (*n*=11), North Jutland (*n*=13), Funen (*n*=3), Zealand (*n*=2), and East Norway/Sweden (*n*=17). Percentages indicate the proportion of genomic variation explained by each PC axis. **(c)** Boxplot of percentage heterozygosity of the highly divergent regions, compared between the sample sites.

**Supplementary Figure 12.**
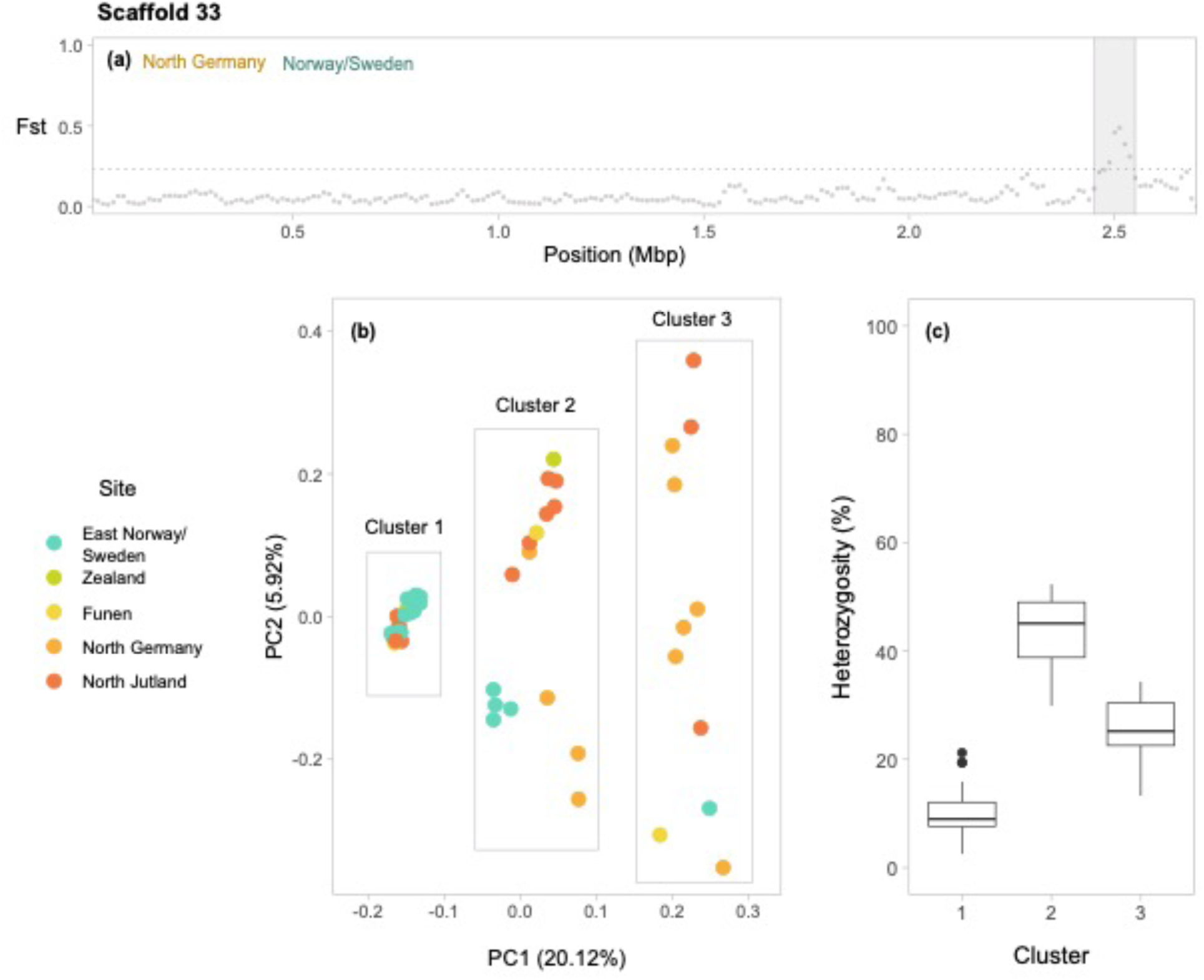
Genomic architecture of the 33rd scaffold of the *B. lapidarius* genome (*n* = 45). The highly divergent region is located between 2.45 and 2.55 Mbp, and contains 134 SNPs**. (a)** Pairwise genetic divergence (Fst) between three *B. lapidarius* populations, measured in sliding windows across the second chromosome with a window size 25kb and a step size of 12.5kb. The populations are North Jutland/Funen (*n*=16), North Germany (*n*=11), and East Norway/Sweden (*n*=17). Coloured labels indicate population pairs. Dotted lines represent the 99^th^ percentile of each Fst frequency distribution. Shaded areas indicate the highly divergent region from which Principal Component Analysis (PCA) and heterozygosity were measured. **(b)** PCA of the highly divergent region. Points represent individual samples. Colours indicate sample site: North Germany (*n*=11), North Jutland (*n*=13), Funen (*n*=3), Zealand (*n*=2), and East Norway/Sweden (*n*=17). Percentages indicate the proportion of genomic variation explained by each PC axis. **c)** Boxplot of percentage heterozygosity of the highly divergent regions, compared between the clusters inferred by Figure S12b.

**Supplementary Figure 13.**
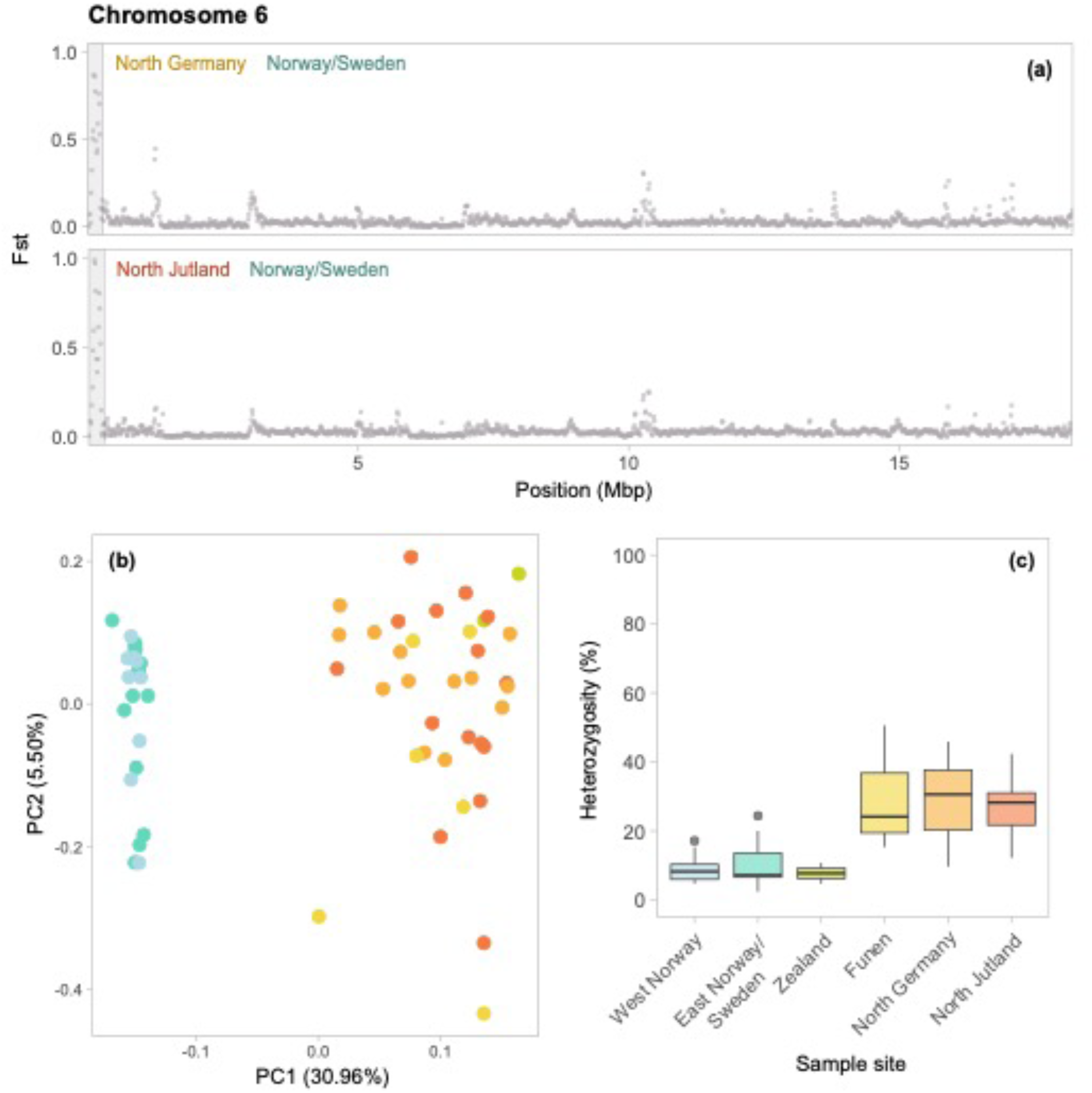
Genomic architecture of the sixth chromosome of the *B. pascuorum* genome (*n* = 61). The highly divergent region is located between 0.09 and 0.26 Mbp, and contains 85 SNPs. **(a)** Pairwise genetic divergence (Fst) between three *B. pascuorum* populations, measured in sliding windows across the second chromosome with a window size 25kb and a step size of 12.5kb. The populations are North Jutland (*n*=15), North Germany (*n*=13), and Norway/Sweden (*n*=25). Coloured labels indicate population pairs. Shaded areas indicate the highly divergent region from which Principal Component Analysis (PCA) and heterozygosity were measured. **(b)** PCA of the highly divergent region. Points represent individual samples. Colours indicate sample site: North Germany (*n*=13), North Jutland (*n*=15), Funen (*n*=6), Zealand (*n*=2), Eastern Norway/Sweden (*n*=15) and Western Norway (*n*=10). Percentages indicate the proportion of genomic variation explained by each PC axis. **(c)** Boxplot of percentage heterozygosity of the highly divergent regions, compared between the sample sites.

**Supplementary Figure 14.**
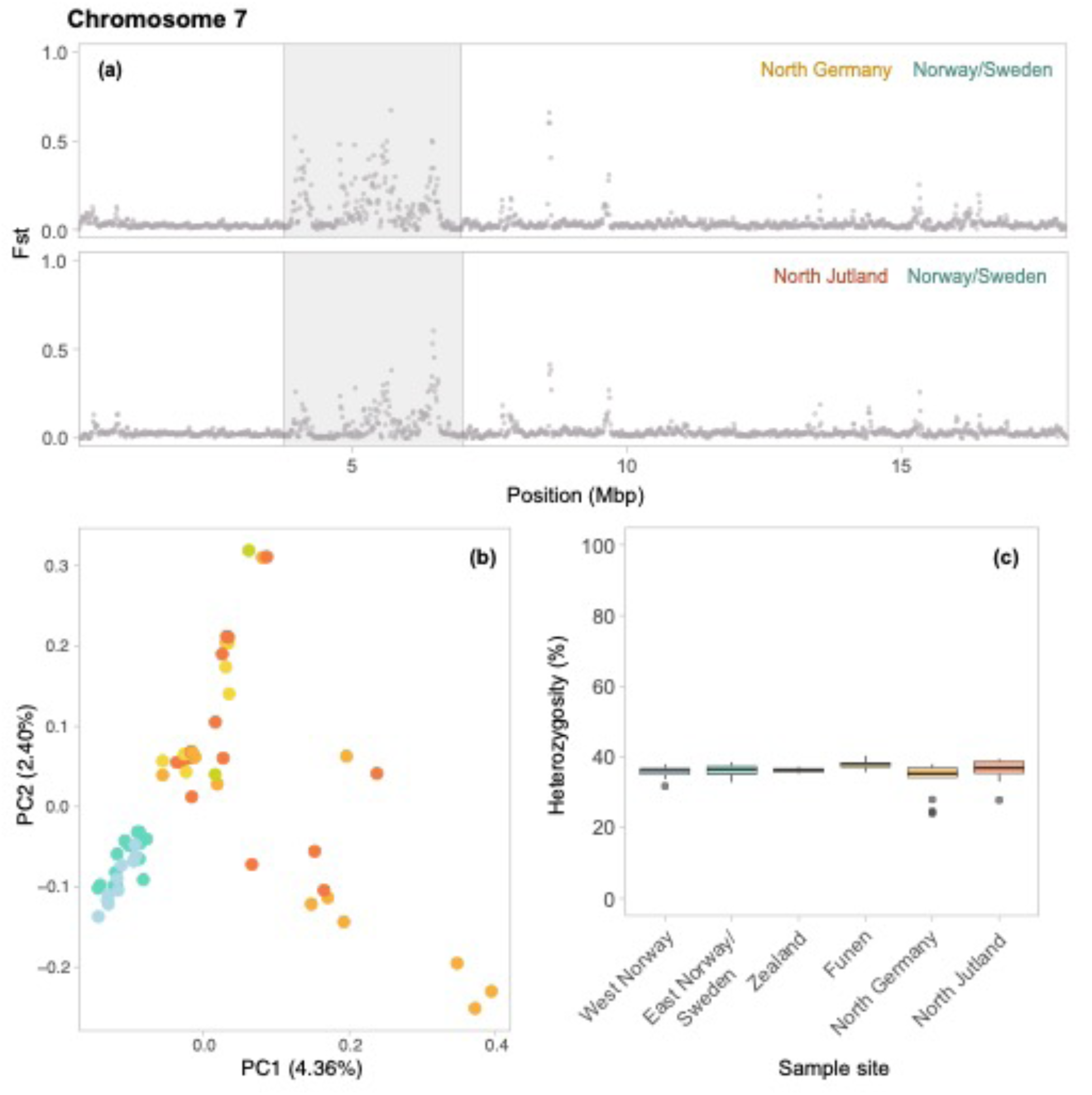
Genomic architecture of the 7th chromosome of the *B. pascuorum* genome (*n* = 61). The highly divergent region is located between 3 and 7 Mbp, and contains 160,284 SNPs. **(a)** Pairwise genetic divergence (Fst) between three *B. pascuorum* populations, measured in sliding windows across the second chromosome with a window size 25kb and a step size of 12.5kb. The populations are North Jutland (*n*=15), North Germany (*n*=13), and Norway/Sweden (*n*=25). Coloured labels indicate population pairs. Shaded areas indicate the highly divergent region from which Principal Component Analysis (PCA) and heterozygosity were measured. **(b)** PCA of the highly divergent region. Points represent individual samples. Colours indicate sample site: North Germany (*n*=13), North Jutland (*n*=15), Funen (*n*=6), Zealand (*n*=2), Eastern Norway/Sweden (*n*=15) and Western Norway (*n*=10). Percentages indicate the proportion of genomic variation explained by each PC axis. **(c)** Boxplot of percentage heterozygosity of the highly divergent regions, compared between the sample sites.

**Supplementary Figure 15.**
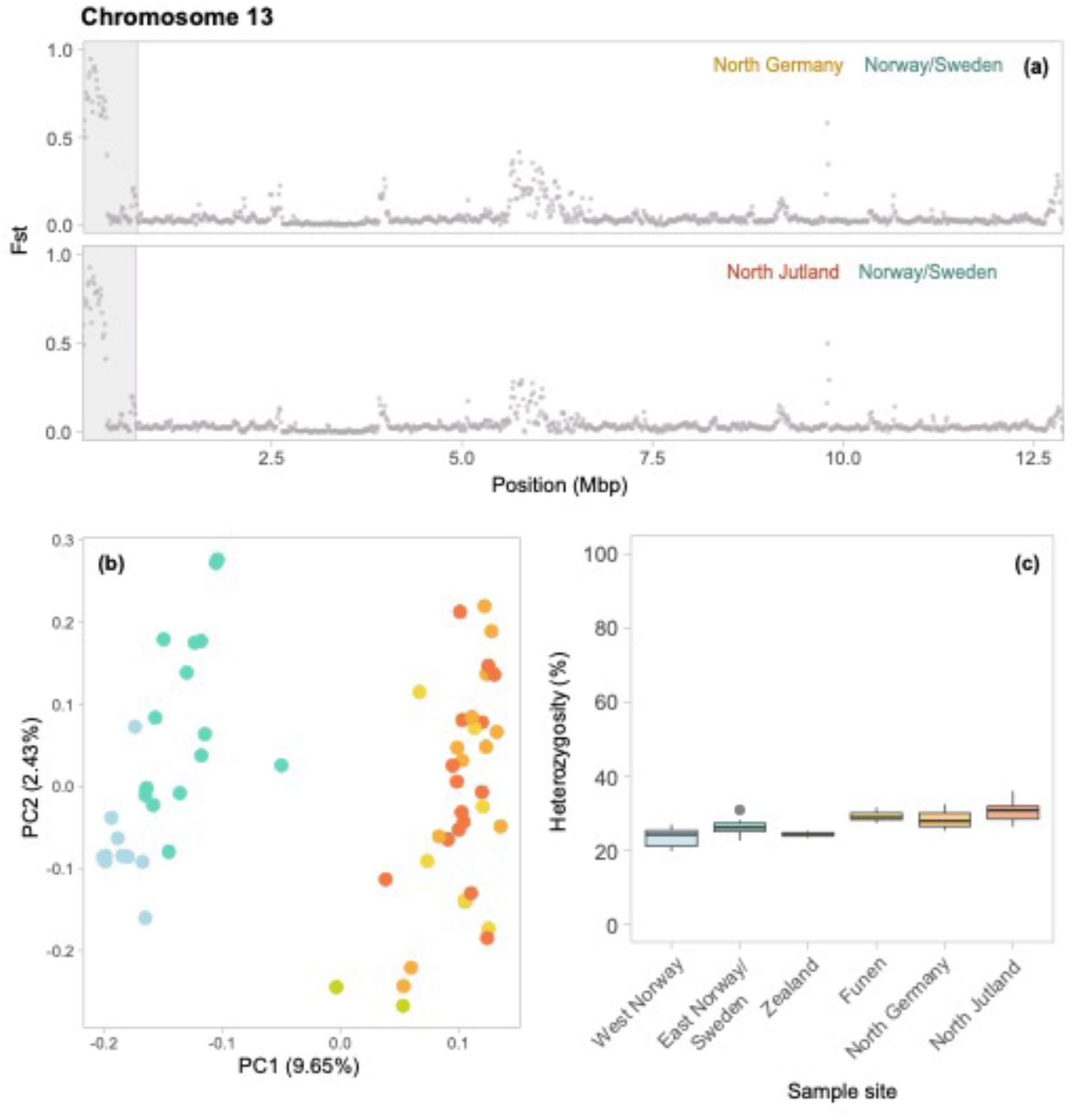
Genomic architecture of the 13th chromosome of the *B. pascuorum* genome (*n* = 61). The highly divergent region is located between 0 and 0.72 Mbp, and contains 2114 SNPs. **(a)** Pairwise genetic divergence (Fst) between three *B. pascuorum* populations, measured in sliding windows across the second chromosome with a window size 25kb and a step size of 12.5kb. The populations are North Jutland (*n*=15), North Germany (*n*=13), and Norway/Sweden (*n*=25). Coloured labels indicate population pairs. Shaded areas indicate the highly divergent region from which Principal Component Analysis (PCA) and heterozygosity were measured. **(b)** PCA of the highly divergent region. Points represent individual samples. Colours indicate sample site: North Germany (*n*=13), North Jutland (*n*=15), Funen (*n*=6), Zealand (*n*=2), Eastern Norway/Sweden (*n*=15) and Western Norway (*n*=10). Percentages indicate the proportion of genomic variation explained by each PC axis. **(c)** Boxplot of percentage heterozygosity of the highly divergent regions, compared between the sample sites.

**Supplementary Figure 16.**
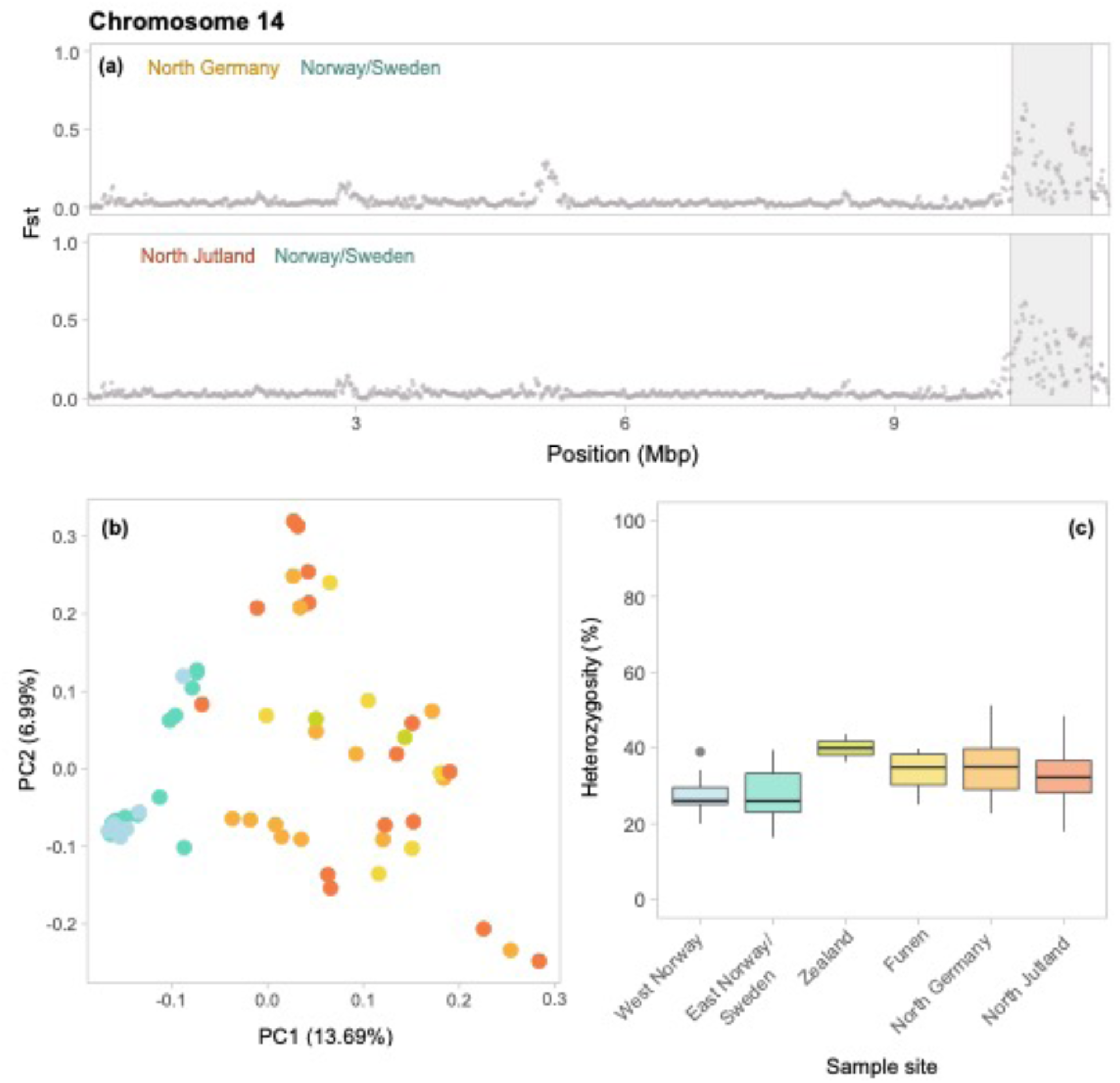
Genomic architecture of the 14th chromosome of the *B. pascuorum* genome (*n* = 61). The highly divergent region is located between 10.3 and 11.2 Mbp, and contains 934 SNPs. **(a)** Pairwise genetic divergence (Fst) between three *B. pascuorum* populations, measured in sliding windows across the second chromosome with a window size 25kb and a step size of 12.5kb. The populations are North Jutland (*n*=15), North Germany (*n*=13), and Norway/Sweden (*n*=25). Coloured labels indicate population pairs. Shaded areas indicate the highly divergent region from which Principal Component Analysis (PCA) and heterozygosity were measured. **(b)** PCA of the highly divergent region. Points represent individual samples. Colours indicate sample site: North Germany (*n*=13), North Jutland (*n*=15), Funen (*n*=6), Zealand (*n*=2), Eastern Norway/Sweden (*n*=15) and Western Norway (*n*=10). Percentages indicate the proportion of genomic variation explained by each PC axis. **(c)** Boxplot of percentage heterozygosity of the highly divergent regions, compared between the sample sites.

**Supplementary Figure 17.**
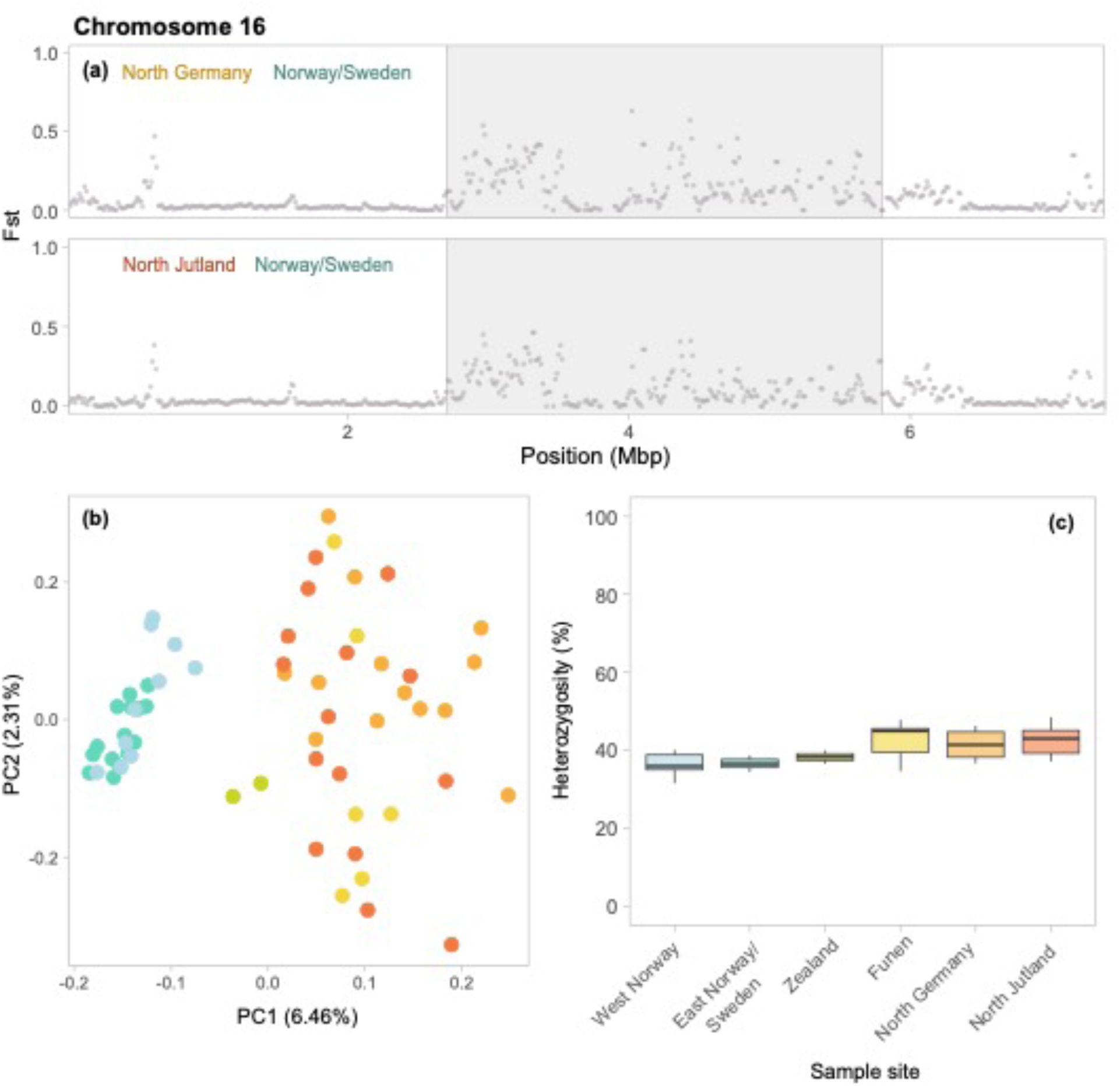
Genomic architecture of the 16th chromosome of the *B. pascuorum* genome (*n* = 61). The highly divergent region is located between 2.7 and 5.8 Mbp, and contains 6672 SNPs. **(a)** Pairwise genetic divergence (Fst) between three *B. pascuorum* populations, measured in sliding windows across the second chromosome with a window size 25kb and a step size of 12.5kb. The populations are North Jutland (*n*=15), North Germany (*n*=13), and Norway/Sweden (*n*=25). Coloured labels indicate population pairs. Shaded areas indicate the highly divergent region from which Principal Component Analysis (PCA) and heterozygosity were measured. **(b)** PCA of the highly divergent region. Points represent individual samples. Colours indicate sample site: North Germany (*n*=13), North Jutland (*n*=15), Funen (*n*=6), Zealand (*n*=2), Eastern Norway/Sweden (*n*=15) and Western Norway (*n*=10). Percentages indicate the proportion of genomic variation explained by each PC axis. **(c)** Boxplot of percentage heterozygosity of the highly divergent regions, compared between the sample sites.

## Supplementary Tables

**Supplementary Table 3.** Per-population mean F statistics for 45 *B. lapidarius* bumblebees.

| **East Norway/**<br>**Sweden** | **Zealand** | **Funen** | **North Germany** | **North Jutland** |
| --- | --- | --- | --- | --- |
| 0.009 | 0.011 | 0.002 | -0.002 | -0.001 |

**Supplementary Table 4.** Per-population mean F statistics for 61 *B. pascuorum* bumblebees.

| **West Norway** | **East Norway/**<br>**Sweden** | **Zealand** | **Funen** | **North Germany** | **North Jutland** |
| --- | --- | --- | --- | --- | --- |
| 0.006 | -0.006 | 0.004 | 0.007 | -0.011 | -0.006 |

